# Regulation of HDAC2-PDX1 by RNF125 defines pancreatic cancer development

**DOI:** 10.1101/2020.01.06.896555

**Authors:** Erez Hasnis, Hyungsoo Kim, Sachin Verma, Yongmei Feng, Ronit Almog, Sivan Matsliah, Vera Vavinskaya, Ron Apelbaum, Offir Ben-Ishay, David Tuveson, Rosalie Sears, Ze’ev A. Ronai

**Author notes:** Correspondence – Ze’ev A Ronai.

## Abstract

There is an urgent need to define mechanisms underlying pancreatic adenocarcinoma (PDA) development. Our studies of ubiquitin ligases that may underlie PDA development led us to identify and characterize RNF125. We show that RNF125 exhibits nuclear expression in acinar cells, with reduced and largely cytosolic expression in ductal cells, PanIN and PDA specimens. We find that RNF125 interacts with histone deacetylase 2 (HDAC2) and promotes its non-canonical K63-linked ubiquitination. Inhibition of HDAC2 activity by RNF125 resulted in elevated expression of the pancreatic and duodenal homeobox 1 (PDX1). Correspondingly, inhibition of RNF125 expression enhanced organoid growth in culture and orthotopic tumor development. Conversely, restoration of PDX1 levels in human or mouse PDA cells and organoids depleted of RNF125, inhibited cell proliferation and growth, while expression of HDAC2 enhanced it. Notably, higher expression of RNF125 and PDX1 coincided with differentiated tumor phenotypes, and better outcome in PDA patients. In demonstrating the importance of RNF125 control of PDX1 expression via HDAC2 ubiquitination in PDA development, our findings highlight markers (RNF125, PDX1) and targets (HDAC2) for monitoring and possible treatment of PDA.

## Introduction

Pancreatic adenocarcinoma (PDA) is notoriously aggressive and resistant to therapy, and is the third leading cause of cancer mortality in the US^1^. Advances in mapping the PDA genomic and transcriptomic landscape have provided insights into deregulated pathways that may underlie this devastating disease. Among the major genetic alterations seen in PDA are mutations in *KRAS, TP53, SMAD4* and *CDKN2A*, implicating hyperactive PI3K, MAPK and WNT signaling, in concert with deregulated cell cycle and DNA damage control mechanisms^2^. Equal attention has been given to epigenetic regulators rewired in PDA, including *DNMT1, DNMT3A, DNMT3B, KDM6A, P300, SMARCA2/4*, several miRs and lnRNAs that impact DNA methylation, histone modifications, and chromatin remodeling^3, 4^. Mapping changes indicative of different phases of PDA development has allowed identification of regulatory cues characteristic PDA development.

Oncogenic pancreatic cells, including PDA, require effective adaptation to low oxygen and low nutrient conditions, as well as the ability to accommodate extracellular matrix (ECM) stiffness and fibrosis^5^. Such adaptation requires rewiring of cellular metabolic and protein homeostasis pathways, resulting in altered metabolic and unfolded protein responses^6–8^. Changes in acinar cell physiology, which are central to pancreatic organogenesis, mark early phases of PDA development. Acinar cells routinely withstand harsh conditions, for which they possess high rate of protein synthesis and secretion^9^, hallamrks of the unfolded protein response (UPR) machinery.

Failure to maintain cellular homeostasis in response to environmental insult such as inflammation or tissue injury, promotes trans-differentiation of acinar cells, known as acinar-to ductal metaplasia (ADM), resulting in a ductal cell lineage^10, 11^. Underlying this process is coordinated regulation of transcriptional and epigenetic networks, including changes in SOX9 and hepatocyte nuclear factor 6 (HNF6) expression, both of which function in somatic stem cell if cellular homeostasis is re-established; however, sustained insult, especially when combined with genetic and epigenetic changes, drives an irreversible state in which ductal cells enter a PanIN phase resulting in a pancreatic intraepithelial neoplasm, a critical step in PDA development^12, 14^. The transcription factor pancreatic and duodenal homeobox 1 (PDX1) is required for differentiation of all pancreatic cell lineages^15^, and PDX1 loss is associated with more aggressive phenotypes of PDA^16^. Another factor implicated in ADM is the nuclear receptor 5A2 (NR5A2), which has been linked with the control of acinar identity and pancreatic inflammation and repair^17, 18^. NR5A2 is more highly expressed in human pancreatic cancer specimens than in normal pancreas and supports proliferation of PDA cell lines^19^.

The ubiquitin system plays a key role in tumor development, progression and resistance mechanisms^20–23^, and changes in activity of several ubiquitin ligases are reportedly seen in PDA development as well as that of other cancers ^21, 24–26^. In search for ubiquitin ligases (UBLs) that are differentially expressed in pancreatic cancer and normal tissues and potentially linked with PDA prognosis we identified the RING-type E3 ubiquitin ligase RNF125. As a 232 aa protein RNF125 harbors a N-terminal RING domain and two central zinc finger domains. Earlier studies have demonstrated that RNF125 controls RIG-I stability, with concomitant effect on the regulation of expression is reduced over the course of acinar-ductal phases and as PDA develops. We identify that RNF125 regulates HDAC2 activity with concomitant effect on the expression of PDX1. Our studies suggest that RNF125-HDAC2-PDX1 axis support early phases in pancreas development, while antagonizes PDA tumor growth,

## Results

### The ubiquitin ligase RNF125 is differentially expressed in the exocrine pancreas

To identify UBLs that contribute to the etiology of PDA, we queried the TCGA database for UBLs differentially expressed in cancer, including melanoma^27, 30^, ovarian cancer^31^, breast cancer^32–34^, colorectal cancer^35, 36^, lung cancer^37^, and head and neck cancer^36, 38^. Of the UBLs that were altered in pancreatic cancer (Fig S1A) we selected UBLs whose expression may be linked with survival. Two RING finger ubiquitin ligases that met these criteria, were *RNF125* and *RNF2.* Whereas *RNF125* expression negatively correlated with survival, *RNF2* expression exhibited an opposite pattern (Fig. S1A).

Decreased RNF125 expression was found in 22% of PDA cases; those changes included reduced transcription (28/149) and partial deletion of the RNF125 locus (5/149) on chromosome 18 (Fig S1A). Independent evaluation of RNF125 expression using the TCGA database confirmed that RNF125 downregulation seen PDA is not common in other tumors (i.e. AML, melanoma; Fig. S1B). Correspondingly, PDA tumor specimens were found to exhibit lower levels of RNF125 expression, compared to that in surrounding normal pancreas (Fig. S1C). Given the differences observed for *RNF125* in PDA we set to further study its possible importance for this tumor type.

We next assessed potential changes in RNF125 expression between the normal pancreas and PDA. Immunostaining of normal pancreas revealed differential expression and localization of RNF125 in the exocrine pancreas. While acinar cells exhibited relatively increased and mostly nuclear RNF125, the ductal epithelium exhibited lower RNF125 levels, and RNF125 was largely excluded from the nucleus in ductal cells (Fig. S2A), implying that RNF125 expression and nuclear localization may be important for the maintenance of normal pancreas. To test this possibility, we assessed pancreatic organization in RNF125 knockout (KO) mice (Figure 1A-B). Notably, smaller pancreata exhibiting decreased expression of developmental (also implicated in acinar) markers (NR5A2, PDX1, GATA6, MIST1; Figure 1C) versus increased ductal markers (PTF1A, SOX9, FGF10; Figure 1D, 1E) were seen in 6-week-old RNF125 KO relative to WT mice. Along these lines, carboxypeptidase A1, amylase and pancreatic lipase, markers associated with acinar cells and exocrine activity were downregulated in pancreata of RNF125 KO mice (Fig. 1F). Chromogranin and insulin levels, associated with endocrine activity, were somewhat lower in RNF125 KO pancreata, although plasma glucose levels were not significantly altered, albeit somewhat increased (Fig. 1G). Correspondingly, pan-cytokeratin staining revealed increased numbers of ductal structures within the pancreatic parenchyma in RNF125 KO mice (Fig. 1H). These observations suggest that RNF125 expression is associated with acinar state as part of normal pancreas homeostasis.

**Fig. 1.**
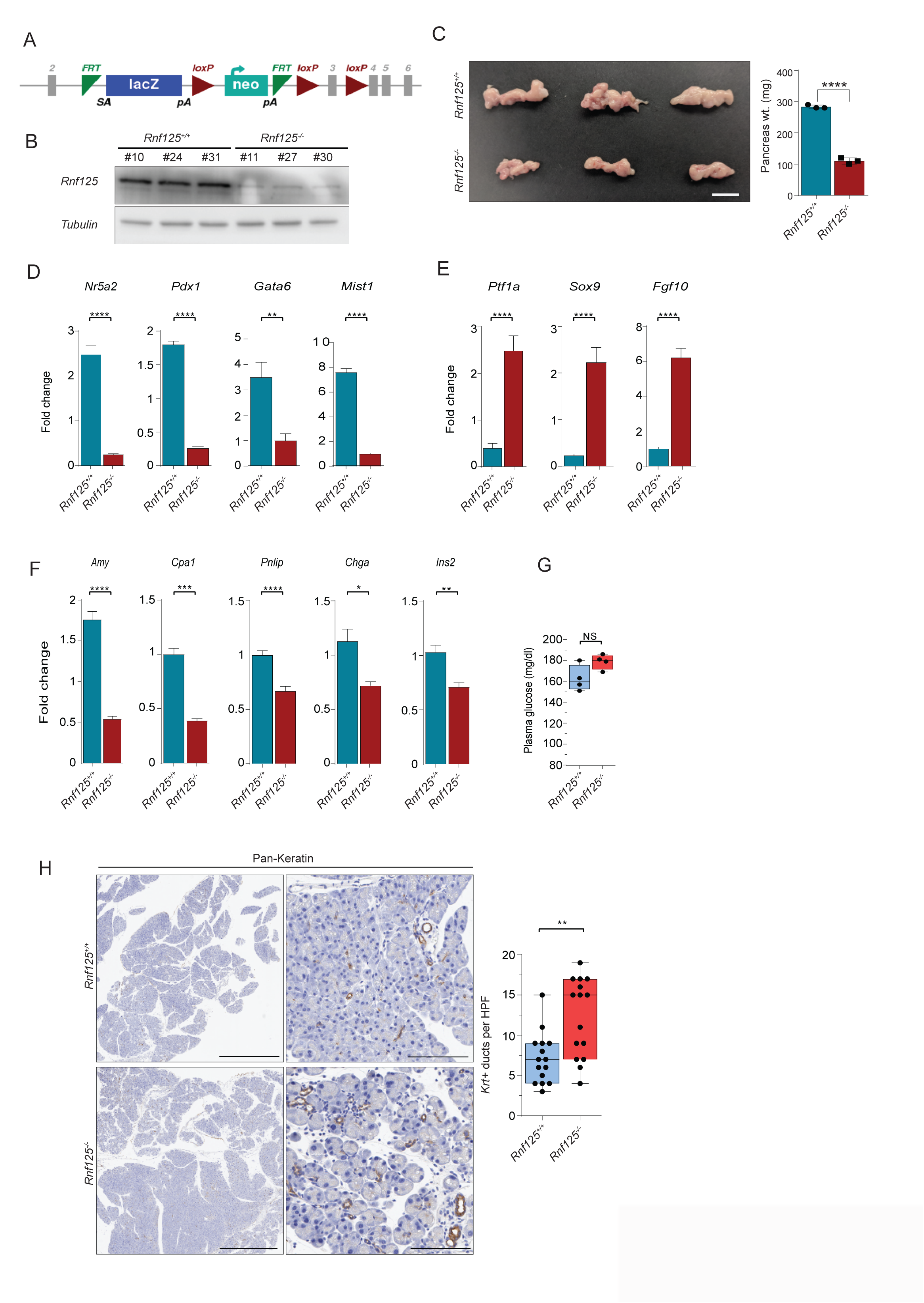
*Rnf125^−/−^* mice exhibit smaller pancreata and impaired acinar differentiation. (**A**) Scheme showing the targeting vector used for *Rnf125* deletion, present in the *Rnf125^−/−^* mice. (**B**) Western blot confirming RNF125 loss in pancreas of *Rnf125^−/−^* mice. (**C**) left, Pictures of pancreata from 6-week-old WT and *Rnf125^−/−^* mice (scale = 1 cm); right, quantification of pancreas weight (n=3, each point represents a different mouse). Student’s t-test was used to calculate statistical significance. (**D**) Relative transcript levels in acinar related genes in pancreata of WT, *Rnf125^−/−^*, or heterozygous mice **(E)** Relative expression of transcripts associated with ductal differentiation in samples outlined in panel D (n=3, results represent means ± SD). Statistical significance was calculated using one-way ANOVA. (**F**) Relative transcript levels of indicated genes functioning in endocrine and/or exocrine function in pancreata of WT, *Rnf125^−/−^* (n=3, results are means ± SEM) (**G**) Fasting plasma glucose levels in indicated mice (n=4, each point represents a different mouse). Statistical significance was calculated using one-way ANOVA. (**H**) left, Pan-Keratin staining of pancreas from indicated genotypes. Left scale bars, 1mm; right scale bars, 100μm. right, Quantification of the number of ductal structures (n=2). HPF = high power field. Student’s t-test was used to calculate statistical significance. \**P*<0.05, \*\**P*<0.01, \*\*\**P*<0.005, \*\*\*\**P*<0.001, NS – nonsignificant.

### Transformed cells exhibit changes in expression and subcellular RNF125 localization

Changes in *RNF125* expression and localization noted in acinar vs. ductal cells (Fig S2A) prompted us to determine the level and localization of RNF125 in a cohort of human PDA samples. While in a few cases (4/22) RNF12*5* expression was high and detectable in the nucleus, most tumor samples (18/22) exhibited relatively low expression of RNF125 that was largely cytosolic (Fig. S2B). RNF125 expression was also reduced in cancer-associated fibroblasts surrounding tumor islets (Fig. S2B). Notably, higher RNF125 protein levels were associated with improved prognosis (Fig. S2B, Table 1). To confirm these observations, we analyzed a TMA consisting of 84 PDA cases. Of those, 32% exhibited low RNF125 levels, 58% were RNF125-negative, and only 10% exhibited strong RNF125 staining, of which only 2 were nuclear (Fig. S2C). Notably, low levels of RNF125 expression coincided with a poorer differentiation grade (Fig. S2C, Table 2). Moreover, lower RNF125 levels in this context were also observed in 8 out of 10 PDA cell lines (Fig. S2D). Overall, decreased nuclear RNF125 was identified in human pancreatic ductal epithelium cells, compared to the mostly acinar normal pancreas, with further decrease in nuclear RNF125 seen in human PDA cell lines (Fig. S2E).

To further assess possible changes in RNF125 expression in tumor development, we monitored RNF125 expression in human pancreatic intraepithelial neoplasm (PanIN), the early precursor of a malignant PDA lesion. Lower levels of RNF125 protein as well as relatively decreased nuclear expression was detected in PanIN samples compared with surrounding acinar cells (Fig. S2F), suggesting that altered expression (lower level and nuclear exclusion) of RNF125 may be linked with PDA development.

To directly assess possible importance of RNF125 in PDA, we assessed the expression and possible function of RNF125 in an established model of pancreatic malignancy. To this end we used an orthotopic PDA derived from the *Pdx1^Cre^;Kras^G12D^;Trp53^R172H^*(KPC) mouse model. KPC-derived tumors exhibited lower levels of RNF125 protein relative to normal mouse pancreas (Fig. S2G). Genetic (shRNA-mediated) inhibition of RNF125 increased the proliferation of three PDA cell lines and enhanced sphere formation by KPC cells (Figs. S3A, S3B), while promoting the growth of KPC-derived organoids (Fig. 2A). Conversely, ectopic expression of the WT but not a RING mutant (RM) form of RNF125 attenuated growth of human PDA-derived lines and reduced the number of nuclei, reflective of proliferation of KPC-driven organoids (Figs. 2A, S3A-B). Ectopic expression of RNF125 also attenuated sphere formation and colony-forming capacity of PDA cell lines (Fig. S3A, C). These data suggest that RNF125 requires its ubiquitin ligase function to exert a tumor growth inhibitory function.

**Fig. 2.**
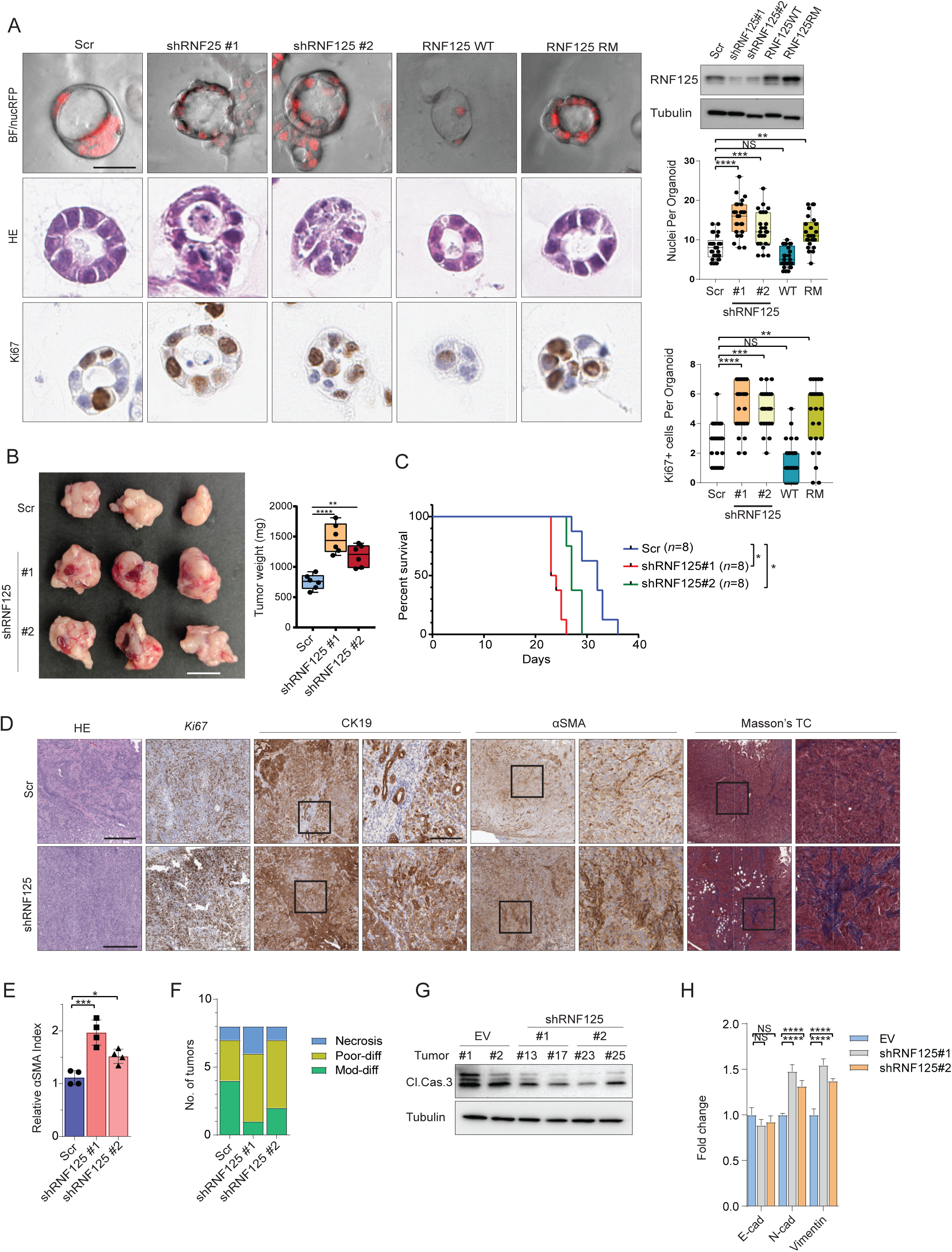
RNF125 attenuates growth of pancreatic cancer organoids and orthotopic tumors. (**A**) left, Representative pictures of 7-day-old KPC organoids in indicated RNF125 genetic backgrounds, in brightfield (upper row, nuclei are RFP-tagged), and stained with H&E (middle row) and Ki67 (lower row). RNF125RM (RNF125 RING finger mutant) right, Western blot confirming RNF125 status (upper), as well as quantification of nuclei (middle) and Ki67-positive nuclei (lower) per organoid (*n*=25 organoids per condition). Scale bar, 100µm. (**B**) left, Representative pictures of orthotopic pancreatic tumors 3 weeks after injection of Scr- or shRNF125-transduced KPC cells. right, Tumor weight (*n*=6 per group). (**C**) Survival of mice harboring KPC orthotopic tumors of indicated genotype. Kaplan-Meier analysis and log-rank test were used, *P* = 0.0324. (**D**) Upper, Analysis of control (Scr) and *Rnf125KD* tumors by indicated staining. Scale bars = 400 μm and 100 μm (insets). **E)** relative αSMA index in representative tumors (n=4) from the indicated groups (Scr, shRNF125#1, shRNF125#2). **(F)** Differentiation grade of orthotopic tumors in the indicated groups (n=8 per group). **(G)** Western blot depicts cleaved caspase 3 levels in representative tumors (n=2) from each group (Scr, shRNF125#1, shRNF125#2). (**H**) Relative expression of indicated transcripts marking the EMT in tumors of different genotype (n=3, results are means ± SD). Statistical significance was calculated using two-way ANOVA. \**P*<0.05, \*\**P*<0.01, \*\*\**P*<0.005, \*\*\*\**P*<0.001, NS – nonsignificant.

Inoculation of KPC-driven organoids stably expressing shRNF125 resulted in larger PDA tumors and a corresponding decrease in mouse survival (Fig. 2B-C). Immunohistochemistry of KPC tumors subjected to RNF125 knockdown revealed increased proliferation, reflected in elevated level of Ki67 expression (Fig. 2D) which was accompanied by elevated alpha smooth muscle actin (αSMA) expression, coupled with increased trichrome staining of collagen, indicative of enhanced fibroblast and desmoplastic involvement (Fig. 2D, 2E). Additionally, inhibition of RNF125 expression reduced the number of malignant ductal structures observed by CK19 staining, indicative of dedifferentiation (Fig. 2D, 2F) and lower level of cleaved caspase 3 (Fig. 2G), reflective of reduced apoptosis. Notably, shRNF125-expressing tumors showed increased expression of EMT markers (Fig. 2H), consistent with appearance of de-differentiation phenotypes (Fig. 2D, 2F). KPC tumors expressing sh*RNF125* did not exhibit changes either T cell infiltration or angiogenesis, reflected by unaltered level of CD45^+^, CD8^+^, and CD31^+^ expression (Fig. S4A). Overall, these observations suggest that RNF125 limits PDA growth.

Given the importance of desmoplastic stroma in PDA development, we monitored possible changes in the expression of stromal markers in tumors subjected to RNF125 KD, compared with WT controls. Among fibrogenesis markers, PDGFRα was significantly increased while TGF-β2, TGF-β3, and TNF-*α* expression were consistently and significantly lower in RNF125 knockdown KPC tumors (Fig. S4B). Noteworthy, the growth of orthotopically implanted KPC tumors in RNF125 KO and WT mice was comparable, and there were no changes in fibroblast activation or tumor grade between genotypes (Fig. S4C-D). These findings suggest that RNF125 contribution to PDA development primarily occurs through its tumor intrinsic effects.

### RNF125 controls PDA growth via differential regulation of PDX1

In light of the changes in RNF125 expression and subcellular localization seen in acinar and ductal pancreatic cells, we asked whether genes implicated in pancreatic lineage differentiation or regeneration may be regulated by RNF125 in PDA. Among genes that were implicated in lineage differentiation and development in the pancreas are the transcription factors PDX1, MNX1, and NR5A2^9^. Indeed, a number of markers of acinar activity, including PDX1 and NR5A2, were upregulated in RNF125^High^ PDA (Fig. 3A). Since NR5A2 and PDX1 function in pancreatic acinar differentiation and are reportedly linked to PDA initiation and growth^19, 40^, we asked whether RNF125 is involved in their regulation also in PDA. RNF125 expression was associated with the expression of both PDX1 and NR5A2 in normal pancreas (Fig. 1D), and inhibition of RNF125 resulted in a lower PDX1 and NR5A2 expression in KPC organoids (Fig. 3B), human PDA cell lines(Fig. S5A) and KPC tumors (Fig. 3C). However, the expression of PDX1, but not NR5A2 was upregulated in RNF125^High^ compared with RNF125^Low^ human PDA (Fig. 3D). These observations suggest that RNF125 expression controls PDX1 but not NR5A2 expression in human PDA, which led us to further focused on the PDX1-RNF125 regulatory axis. Indeed, ectopic expression of WT, but not RING mutant (RM), form of RNF125 restored PDX1 expression, suggesting that for its control of PDX1 transcription RNF125 requires its ubiquitin ligase function (Fig 3E). Notably, analysis of a large patient cohort consisting of four different pancreatic cancer subtypes, low expression of RNF125 and PDX1 was seen in squamous type PDA, which is associated with low survival^16^ (Fig. 3F).

**Fig. 3.**
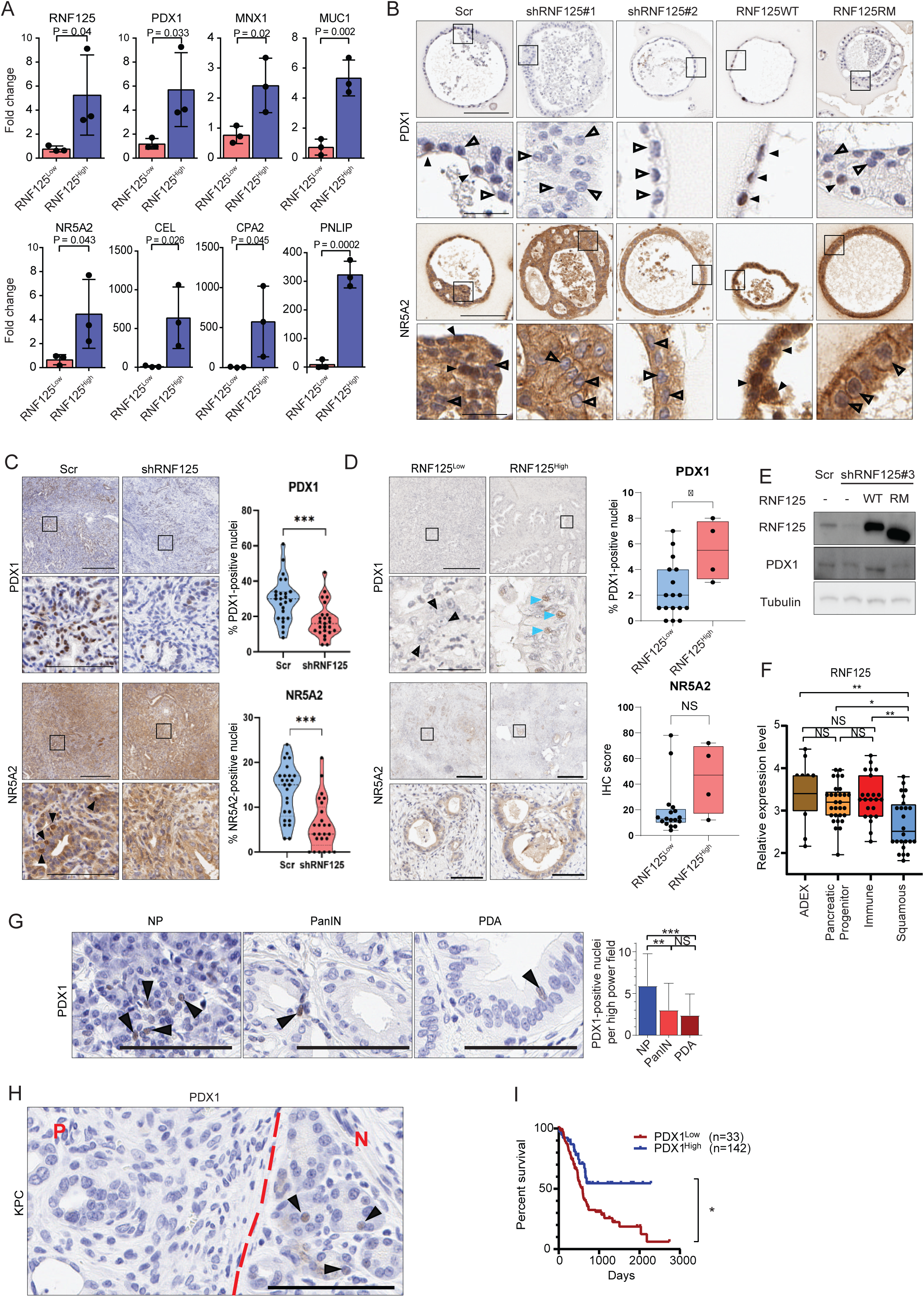
RNF125 regulates PDX1 and NR5A2 transcript levels in pancreatic cancer cells. (**A**) Quantification of indicated genes in RNF125^Low^ vs RNF125^High^ patient samples. Statistical analysis by single-tailed Student’s t-test(*n* = 3 per group). (**B**) IHC staining of indicated proteins in 7-day-old KPC organoids of control (Scr), shRNF125 (#1 or #2) or RNF125RM (RING finger mutant); Scale bars = 300 μm (right), 50 μm (insets). Arrowheads indicate positive (filled) or negative (empty) nuclei stain. (**C**) Left, IHC staining with indicated proteins of KPC orthotopic tumors. Scale bars = 400 μm (upper panels) and 100 μm (insets). Right, Quantification of results from 25 fields from 2 representative mice per group. Student’s t-test was used to calculate statistical significance. \*\*\**P*<0.005. (**D**) Left, IHC staining with indicated markers in RNF125^Low^ (*n*=17) vs. RNF125^High^ (*n*=4) human PDA samples. PDX1-negative (black arrowheads) and - positive (blue arrowheads) nuclei are marked. Scale bars = 400 μm (upper panels) and 75 μm (insets). Student’s t-test was used to calculate statistical significance. \**P*<0.05. (**E**) Western blot analysis with indicated antibodies in Mia PaCa-2 cells co-transfected with shRNF125 plasmid, which targets the 3’UTR region, and either wild type (WT) or RING mutant (RM) forms of RNF125, both of which lack that region. (F) RNF125 expression in four different pancreatic cancer subtypes based on^19^. Statistical significance was calculated using one-way ANOVA. (**G**) Right, representative PDX1 staining in normal human pancreas (NP) and in PanIN and PDA specimens. Black arrowheads indicate PDX1-positive nuclei. Scale 100 μM. Left, quantification of staining (n=3; 10 fields counted for each) (**H**) Representative PDX1 staining in pancreatic samples from 3-month-old KPC mice, showing normal pancreas (N) and PanIN (P). Scale = 50 μM (PDX1).(**I**) Kaplan-Meier plot comparing PDA patients with PDX1^High^ versus PDX1^Low^ tumors; two-sided log-rank *P* = 0.0069. \**P*<0.05, \*\**P*<0.01, \*\*\**P*<0.005, \*\*\*\**P*<0.001, NS – nonsignificant.

To further assess the status of RNF125 and PDX1 in PDA, we monitored changes in the expression of these proteins at different stages of pancreatic cancer development in human samples. The level of PDX1 expression was somewhat lower in PanIN and PDA compared with the normal pancreas (Fig 3G). Likewise, PanIN from the pancreas of the KPC mice exhibited limited changes in PDX1 expression (Fig. 3H). These observations point to direct correlation between PDX1 and RNF125 expression.

Analysis of PDA patient specimens in the TCGA dataset revealed that high PDX1 expression is associated with prolonged survival (Fig. 3I). Notably, tumors expressing both RNF125 and PDX1 showed better survival, compared with tumors expressing low PDX1 or low PDX1/RNF125 (Fig. S5B, S5C)

To assess the potential effect of RNF5-PDX1 expression on PDA we asked whether PDX1 mediates RNF125 effects. On its own, altered PDX1 expression elicited a notable effect on pancreatic tumor growth based on analysis of 2D growth of PDA cell lines (Fig. S5D and organoids (Fig. 4A). Notably, while inhibition of PDX1 expression using siRNA did not led to a marked increase in PDA cell line proliferation, ectopic expression of PDX1 was able to significantly inhibit PDA cell proliferation (Figs. 4A, S5D). We then injected KPC organoids orthotopically into syngeneic C57BL/6 mice and monitored the formation of pancreatic tumors (Fig 4B). As expected, inhibition of RNF125 expression in organoids further accelerated tumor growth *in vivo*. Notably, overexpression of PDX1, attenuated the enhanced growth seen in RNF125 knockdown organoids and resulted in smaller tumors that exhibited decreased proliferation, and increased apoptosis (Fig. 4C). Notably, loss of PDX1 in MiaPaCa-2 cells, increased epithelial to mesenchymal transition (EMT), which was monitored using N-Cadherin and Vimentin as markers (Fig. S5E). Consistent with these observations, overexpression of PDX1 reverted the increased EMT, seen upon RNF125 KD (Fig. S5F). These findings substantiate the importance of RNF125 and PDX1 in PDA development/progression.

**Fig. 4.**
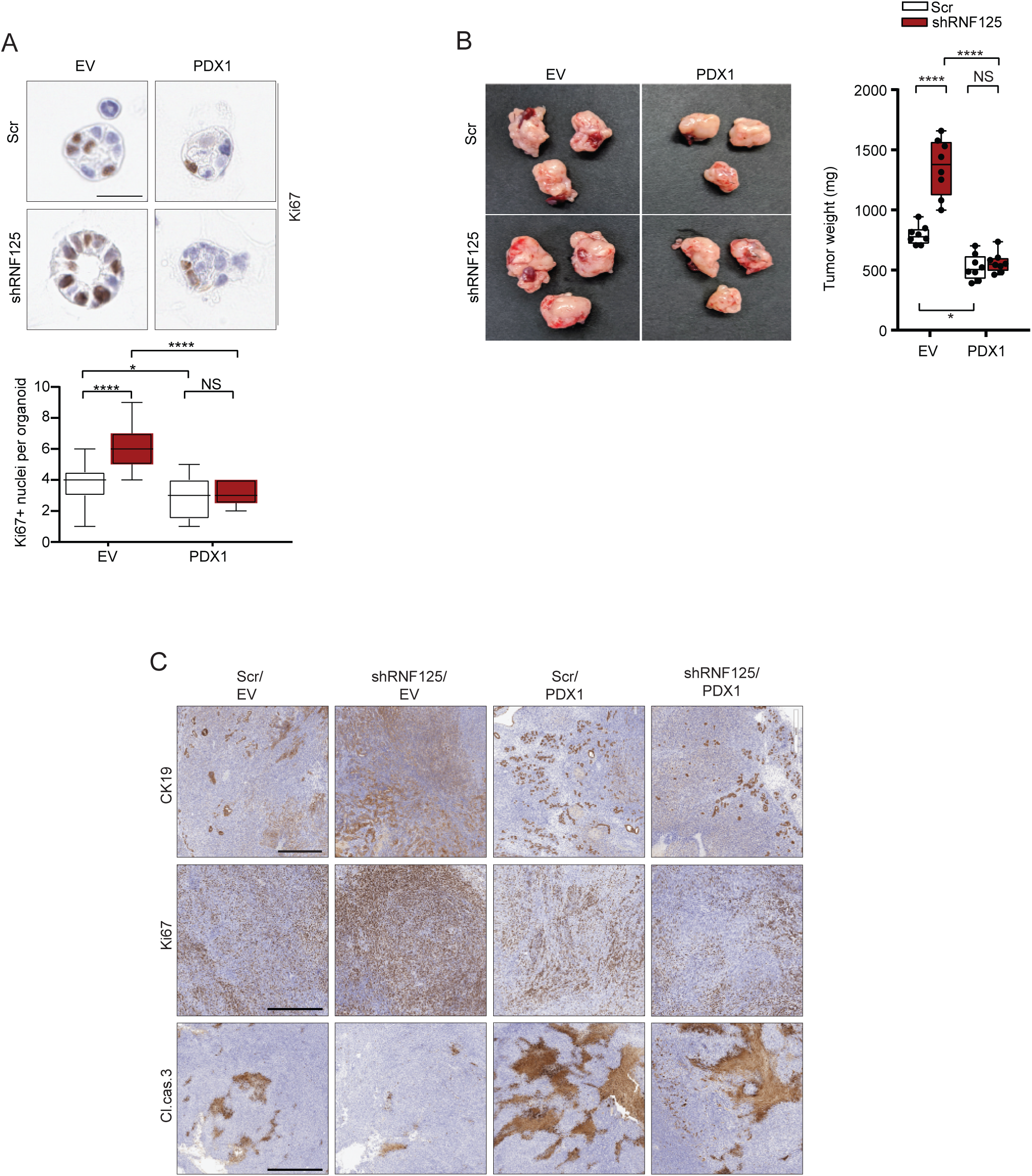
PDX1 but not NR5A2 rescues RNF125 knockdown phenotypes in PDA cells. (**A**) top, Representative Ki67 staining in 48-hour-old KPC organoids transduced with shContol (upper row) or shRNF125 (lower row), and overexpressing empty vector (EV; left), or PDX1 (right). Scale bar, 50μm. Bottom, number of Ki67-positive nuclei per organoid in corresponding conditions (**B**) left, Representative KPC orthotopic tumors, 3 weeks after injection of cells corresponding to those shown in (A). right, Tumor weights in corresponding conditions (*n*=8 mice per group). (**C**) IHC analysis of tumors corresponding to those shown in panel B with indicated staining.

### Nuclear RNF125 interacts with histone deacetylase 2 to decrease its repressive function and attenuate tumor growth

As a ubiquitin ligase, RNF125 should limit either stability or activity of its putative substrates. Yet, given that RNF125 altered PDX1 mRNA levels (Fig S5A), we sought to identify a regulatory protein intermediate as a potential RNF125 substrate. To do so, we performed mass-spectroscopy (MS) using RNF125 as bait, which led us to identify both nuclear and cytosolic RNF125-interacting proteins, some previously reported to play a role in PDA (Fig 5A, Table S1). Given that nuclear RNF125 expression is more prominent in nuclei of non-transformed cells and that PDX1 transcription was linked with RNF125 activities, we searched for putative substrates that govern gene transcription, and thus focused on nuclear RNF125-interacting proteins. Our criteria for selection of substrates included: (i) the ability of an RNF125 substrate to alter transcription of PDX1, and (ii) inhibition of the substrate should rescue phenotypes seen following RNF125 inhibition. When we tested the nuclear RNF125-interacting proteins based on these criteria, only HDAC2 inhibition effectively restored the level of PDX1 expression, which were inhibited upon RNF125 knockdown (Fig. 5B). Immunoprecipitation (IP) of ectopically expressed RNF125 in the MiaPaCa-2 cell line confirmed HDAC2 interaction (Fig. 5C), and further subcellular fractionation analysis of endogenous proteins showed that interaction was also detected in the nucleus (Fig. 5D-E). Notably, the degree of RNF125/HDAC2 interaction and HDAC2 levels were unaltered upon treatment of cells with proteasome inhibitors (Fig. 5D), suggesting that RNF125 does not alter HDAC2 stability.

**Fig. 5.**
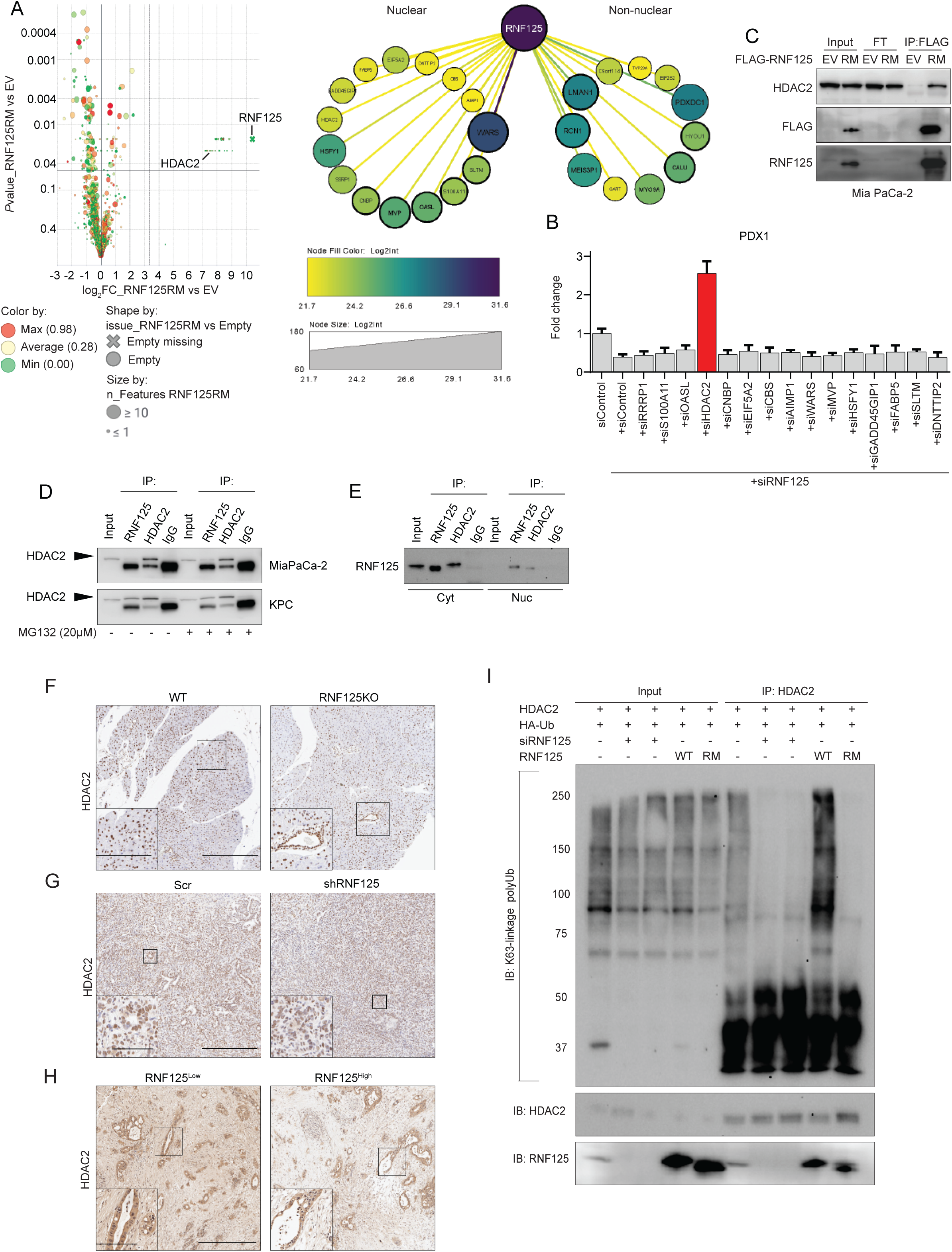
HDAC2 is an RNF125 substrate. (**A**) left, LC-MS/MS analysis of cultured Mia PaCa-2 cells using RNF125RM as bait. right, Networks of nuclear and non-nuclear RNF125 interactors. (**B**) PDX1 transcript levels in Mia PaCa-2 cells transfected with indicated siRNAs. Results are means ± SEM in a representative experiment. (**C**) Validation of RNF125-HDAC2 interaction based on western analysis with the indicated antibodies in total lysates, flow-through (FT) and the FLAG-immunoprecipitated fraction of Mia PaCa-2 cells expressing empty vector (EV) or FLAG-tagged RNF125^RM^. (**D**) Interaction of endogenous RNF125 with HDAC2 in Mia PaCa-2 and KPC cells based on western blotting with an HDAC2 antibody in input and indicated immunoprecipitation fractions, and with or without the proteasome inhibitor MG132. (**E**) Western blot of RNF125 in input and indicated immunoprecipitated fractions showing interaction of endogenous RNF125 with HDAC2 in the cytoplasm (Cyt) and nucleus (Nuc) of Mia PaCa-2 cells. (**F)** HDAC2 staining of normal pancreas from WT and *Rnf125^−/−^* mice. Scale bar in low magnification panels, 250μm and in inset, 100 μm. (**G**) HDAC2 staining of control and shRNF125 KPC tumors. Scale bar in low magnification panels, 400μm and in inset, 50 μm. (**H**) HDAC2 staining in RNF125^Low^ vs. RNF125^High^ human PDA. Scale bar in low magnification panels, 600 μm and in inset, 150μm. (**I**) K63 HDAC2 ubiquitination of as analyzed in HEK293T cells overexpressing HDAC2, HA-Ub, and different RNF125 levels. IP, HDAC2; immunoblot, anti K63-linked Ub antibody.

Indeed, levels of HDAC2 protein in pancreas were comparable in RNF125 WT and KO mice (Fig. 5F). Likewise, both WT and RNF125 knockdown KPC tumors exhibited similar HDAC2 levels (Fig. 5G). Furthermore, human PDA specimens expressing high RNF125 levels exhibited levels of HDAC2 comparable to specimens expressing low RNF125 levels (Fig. 5H). Lastly, inhibition of RNF125 expression in PDA cell lines did not alter basal levels or half-life of HDAC2 (Fig. S6A-B). Collectively, these observations suggest that RNF125 does not affect HDAC2 stability.

Next, we assessed potential effects of RNF125 on HDAC2 ubiquitination. Indeed, inhibition of RNF125 expression in HEK293T cells over-expressing HDAC2 decreased levels of ubiquitin chains of the K63-topology on HDAC2, as compared to WT controls (Fig. 5I). Conversely, ectopic RNF125 expression increased the level of HDAC2 K63-linked ubiquitinated HDAC2 relative to controls, whereas overexpression of the RNF125 RING mutant inhibited HDAC2 non-canonical ubiquitination (Fig. 5I). Likewise, RNF125 inhibition reduced the degree of endogenous HDAC2 K63-linked ubiquitination in both the human MiaPaCa-2 and mouse KPC cells (Fig. S6C-D), while no changes were detected in the levels of HDAC2 K48-ubiquitination (Fig. S6E). Expression of a K63R mutant form of ubiquitin, which should selectively block K63-linked ubiquitin chain, decreased the degree of RNF125-dependent HDAC2 ubiquitination (Fig. S6F).

Given that HDAC2 reportedly represses transcription^41^, and since HDAC2 was subjected to K63 ubiquitination, we assessed effects of HDAC2 on PDX1 expression. Inhibition of HDAC2 expression with shRNA in KPC organoids led to increased PDX1 expression, which was unchanged following RNF125 inhibition (Fig. 6A). Similarly, HDAC2 knockdown in human PDA cells also increased PDX1 protein levels concomitant with increased histone H3K27 acetylation (Fig. S7A). Chromatin IP analysis also revealed enrichment of HDAC2 on the PDX1 promoter, which coincided with lower levels of H3K9ac and H3K27ac on PDX1 promoter (Fig. 6B-C).

**Fig. 6.**
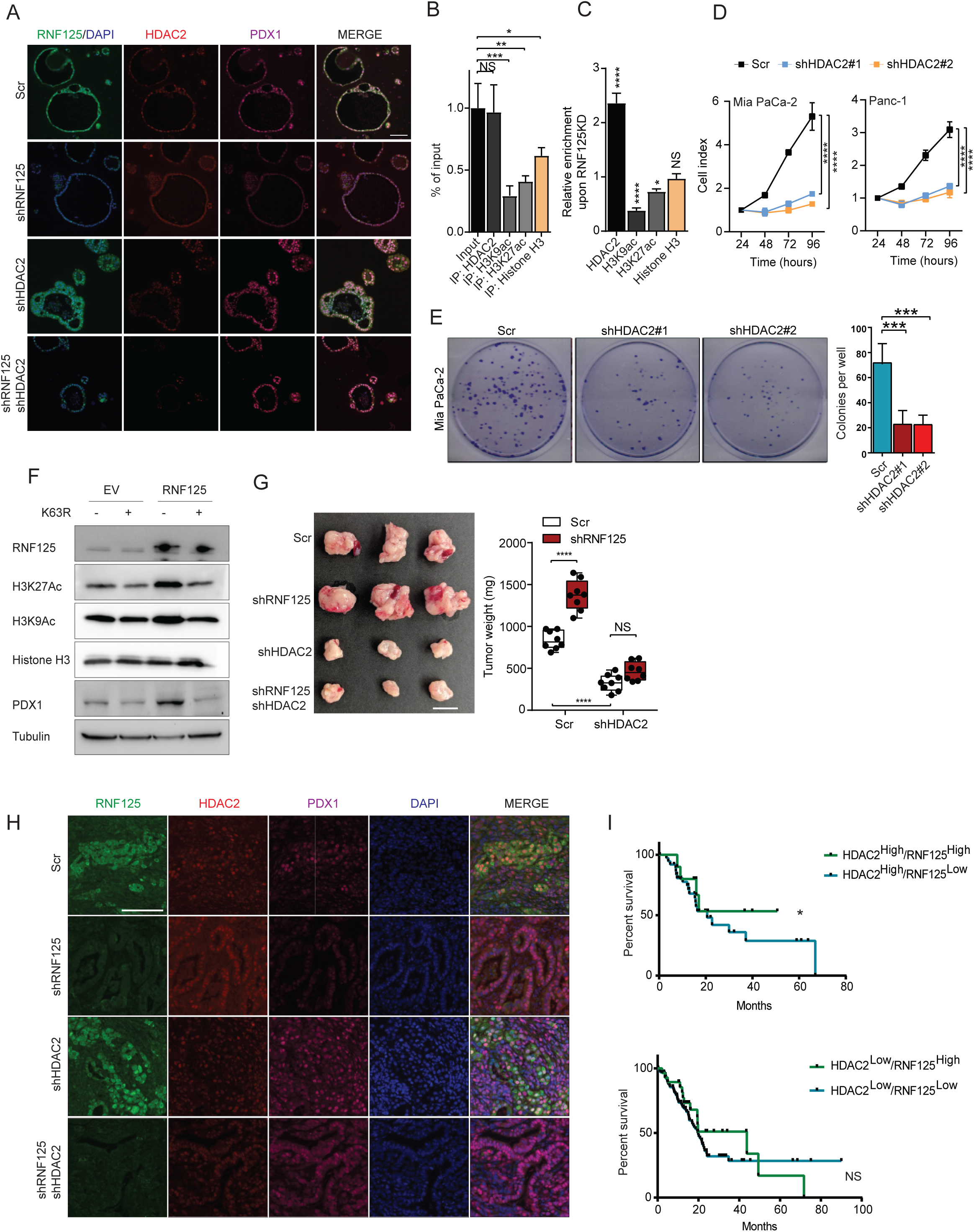
HDAC2 regulates PDX1 levels in pancreatic cancer. (**A**) Immunofluorescence analysis of 7-day-old WT, single knockdown, or double knockdown KPC organoids probed with indicated antibodies. Scale bar, 100 μm. (**B**) Relative levels of indicated transcripts on PDX1 promoter (shown as % of input), in the ChIP performed in KPC cells. Shown are results represents 2 experiments. (**C**) ChIP analysis reveals relative enrichment of HDAC2 on the PDX1 promoter following RNF125 knockdown in KPC cells. Shown are representative data from 2 experiments. (**D**) Proliferation of indicated cell lines shown after transfection with indicated plasmids. One-way ANOVA was used to calculate statistical significance at the 96-hour time point. (**E**) Anchorage-dependent colony-forming assay of Mia PaCa2 cells after transfection with Scr control or two different shHDAC2 plasmids. (**F**) Western blot analysis of indicated proteins in Mia PaCa-2 cells expressing empty vector (EV) or RNF125 in the presence or absence of the K63R Ub mutant. (**G**) Pancreatic tumors 3 weeks after orthotopic injection of Scr-, shRNF125-, shHDAC2-, or shRNF125/shHDAC2-transduced KPC cells. *n*=8. Two-way ANOVA was used to calculate statistical significance. (**H**) RNF125, PDX1, or HDAC2 staining in tumors shown in (G). (I) Kaplan-Meier plots for HDAC2^High^ (upper) and HDAC2^Low^ (lower) PDA patients based on RNF125 levels. \**P*<0.05, \*\**P*<0.01, \*\*\**P*<0.005, \*\*\*\**P*<0.001, NS – nonsignificant.

Notably, attenuated HDAC2 expression by itself significantly inhibited the proliferation of PDA cell lines and organoids (Figs. 6D-E). In addition, in organoid cultures, reduced HDAC2 rescued changes seen after RNF125 knockdown, namely, increased organoid proliferation a (Fig. S7B). Furthermore, ectopic expression of K63R mutant ubiquitin, which attenuates the formation of K63 ubiquitin chains, restored the degree of H3K27 acetylation and blocked increases in PDX1 expression seen following RNF125 expression (Fig. 6F). Correspondingly, orthotopic injection of KPC organoids subjected to HDAC2 knockdown resulted in smaller and more differentiated tumors relative to control tumors (Fig. 6G). Notably, HDAC2 knockdown tumors also showed increased PDX1 expression independent of RNF125 manipulation, confirming the role of HDAC2 in the regulation of PDX1 expression (Figs. 6G, 6H, S7C). High expression of HDAC2 and RNF125 in PDA specimens also correlated with improved patient survival, an effect not seen when only HDAC2 was queried, further suggesting that HDAC2 is required for RNF125-related outcomes (Fig. 6I).

## Discussion

Understanding mechanisms that underlie different phases of PDA development and progression is key to developing novel treatment modalities for this therapy-resistant tumor. Here, we identified a role for the ubiquitin ligase RNF125 in normal pancreas and in PDA development.

Relatively higher expression of RNF125 was seen in acinar cells, suggesting that this ubiquitin ligase functions in normal pancreas development or homeostasis. During pancreas transformation and PDA, RNF125 expression was reduced and was predominantly cytosolic, suggesting that pancreatic transformation either requires reduced RNF125 expression, or its nuclear exclusion, or both. Finding that RNF125 loses its nuclear function during PDA formation is consistent with the observation that RNF125 inhibition accelerates growth of PDA cell lines in culture and growth of PDA *in vivo*. Nuclear exclusion of RNF125 is likely maintained by post-translational modifications during the course of PDA formation and may linked with the reduced level of RNF125 expression.

Our studies further demonstrate that RNF125 indirectly regulates expression of the PDA-associated transcription factor PDX1. Evidence supporting the importance of RNF125 and its regulation of PDX1 for PDA development comes from our rescue studies showing that PDX1 overexpression restores phenotypes seen upon RNF125 knockdown, both in organoids and in orthotopic tumor models.

Notably, our studies identified HDAC2 as an RNF125 substrate, and suggest that HDAC2 activity is regulated by RNF125-mediated K63-linked ubiquitination. Ubiquitination by RNF125 altered HDAC2 activity, as reflected by histone acetylation of the PDX1 promoter region, an activity that may have also been impacted by reduced HDAC2 recruitment to that promoter. Notably, we confirmed a link between HDAC2 and PDX1 in series of studies performed in both KPC organoids and in orthotopic tumor xenografts, in which HDAC2 inhibition rescued growth phenotypes observed following RNF125 inhibition. RNF125 control of HDAC2 activity likely regulates transcription of genes other than PDX1, consistent with our finding of altered transcription of other pancreatic differentiation factors in RNF125 KO mice. Accordingly, better prognosis seen in patients whose PDA specimens harbor elevated RNF125 and HDAC2 expression suggests that components of a transcriptional network along this regulatory axis could serve as markers for PDA development and in stratifying patients for treatment with HDACi.

Relatively lower RNF125 levels were observed in PDA compared to healthy tissues, as was an inverse correlation between RNF125 expression and differentiation grade of human tumors and patient survival. Correspondingly, reducing RNF125 expression increased tumor growth, a phenotype significantly attenuated after either PDX1 overexpression or, more effectively, HDAC2 knockdown. Thus, our work reveals a mechanism controlling PDX1 suppression by HDACs, that may underlie dedifferentiation and cell transformation. Would HDACi offer a novel therapeutic modality for PDA? Our findings support such a possibility, which should be further confirmed experimentally. For one, potential use of domatinostat, a class I HDAC inhibitor, which is currently being evaluated against microsatellite stable colorectal cancer in clinical trials (<u>NCT03812796</u>), could also be evaluated in RNF125 low-expressing PDAs.

In summary, our data suggest that HDAC2 inhibition by RNF125 plays a critical role in maintaining acinar differentiation in the exocrine pancreas. We conclude that RNF125 deregulation is important driver of pancreatic transformation, through its control of HDAC2 activity and transcriptional regulation of key factors in PDA, including PDX1.

## Acknowledgements

We thank Colin Goding and Serge Fuchs for their valuable comments and members of the Ronai lab for advice and continued discussion. This study was supported by TRDRP grant T30IP0941 (to ZR) and NCI grant R35CA197465 (to ZR). Support through grant P30 CA030199 to the analytical genomics, vivarium, microscopy, and histology core facilities at Sanford Burnham Prebys Medical Discovery Institute NCI Cancer Center is gratefully acknowledged

## Materials and Methods

### Human specimens and pancreatic cancer tissue microarray

All human experiments were approved by the institutional review board (0041-17-RMB) at the Rambam Medical Center, Haifa, Israel. Patients provided written informed consent and were de-identified. Pancreatic cancer tissue samples (N=33) with paired adjacent normal pancreas were obtained from metastasis free patients undergoing pancreatic resection. All clinical cases were reviewed by an expert pancreatic oncologist (R.A.) who selected cases based on sufficient clinical data, accurate determination of recurrence time, and complete follow-up. Material received was frozen in −80°C for protein and RNA analysis, while formalin-fixed tissue was used for immunohistochemistry. Confirmation of tumor or normal tissue, and of differentiation grade, was based on pathological assessment using H&E-stained formalin-fixed tissue. Stained slides were interpreted by a dedicated pancreatic pathologist, and only samples verified to have clear PDA morphology were further evaluated. Clinical-pathological data accompanying the samples included age, sex, clinical stage, differentiation grade, and overall survival (Table 1). A pancreatic cancer tissue microarray (TMA) was purchased from Biomax US. Clinical-pathological data accompanying TMA samples included age, sex, clinical stage, and differentiation grade (Table 2).

### Animal studies

All animal experiments were reviewed and approved by the SBP IACUC (AUC 18-001). Heterozygous RNF125KO mice were purchased from Eucomm. Wild-type C57BL/6J male mice were purchased from the Jackson Laboratory. For animal studies, male mice of similar age (6 to 8 weeks) were used. For orthotopic injection of KPC cells, shaved mice were anesthetized using continuous isoflurane, and their abdomen was sterilized. An upper left laparotomy (5-10 mm) exposed the peritoneal cavity, allowing exposure of the pancreas for injection of KPC organoids (which originated from a single matrigel dome containing 1×10^5^ cells suspended in 30 μl sterile PBS). Successful injection was confirmed by formation of a liquid bleb at the injection site with no fluid leakage. The pancreas was then placed back into the peritoneal cavity. The peritoneum was sutured by a single 5-0 silk suture, and skin was closed by clipping. After mice were sacrificed, tumors were removed and frozen for protein and RNA extraction or fixed in formalin (for immunohistochemistry and immunofluorescence).

### Cell lines

The human pancreatic cancer cell line Mia PaCa-2 was kindly provided by the G. Powis laboratory (SBP), and T3M4 was kindly provided by the M. Korc laboratory (UI). Panc-1, BxPC-3, AsPC-1, HPAF-II, CFPAC-1, Capan-1, Capan-2 and human pancreatic ductal epithelium (HPDE) cells were kindly provided by the A. Lowy group (UC San Diego). Cells derived from a primary pancreatic KPC tumor (*Kras^+/LSL-G12D^;Trp53*^+/LSL-R172H^*;Pdx-Cre)* were generated by the D. Tuveson group (CSHL) and maintained as orthotopically-injected tumors^42^. Human PDA lines were cultured as recommended by ATCC. HPDE were cultured in keratinocyte serum free (KSF) medium supplemented with EGF. KPC cells were cultured in Dulbecco’s modified eagle medium (DMEM) supplemented with fetal bovine serum (FBS, 10% v/v), penicillin and streptomycin.

### Organoid culture

Organoids were prepared as described^43^. Briefly, bulk orthotopic KPC tumors were minced and digested overnight with collagenase XI and dispase, washed, counted, embedded in growth factor-reduced (GFR) matrigel (Corning, 1 × 10^5^ cells suspended in a 50μl matrigel dome), and cultured in murine complete medium [Advanced DMEM/F12 medium supplemented with HEPES (1x, Corning), Glutamax (1x, Gibco), penicillin/streptomycin (1x, Gibco), B27 (1x, Invitrogen), N-acetyl-L-cystenine (1mM, Sigma), A83-01 (500 nM, Sigma), RSPO-1 conditioned medium (10% v/v, Cultrex® Rspo1 produced by HEK293T cells, <u>Trevigen),</u> mNoggin (0.1 μg/ml, Peprotech), mouse epidermal growth factor (mEGF, 50 ng/ml), Gastrin-I (10nM, Peprotech), human fibroblast growth factor 10 (hFGF10, 100 nm/ml, Peprotech), and nicotinamide (10 mM, Sigma)]. To isolate protein or RNA, or for orthotopic injection of organoids, matrigel was manually disrupted in cold PBS at indicated time points, and organoids were washed and collected. For organoid immunofluorescence and immunohistochemistry, the matrigel dome was fixed in formalin, embedded and cut.

### Cycloheximide chase assay

Cycloheximide chase was performed as previously described^44^. Briefly, cycloheximide (50 μg/ml) was added to cells for indicated times, and cell lysates were analyzed with indicated antibodies.

### Cell viability assay in 2D culture

Cell growth was assayed using the standard trypan blue exclusion assay or ATPlite (Perkin Elmer). Briefly, 4 hours after transfection, PDA cells were washed, harvested, and cultured in 96-well plates with a transparent bottom (5,000 cells/well). For paclitaxel treatment, drug was added 24 hours after seeding, and then cells were incubated 48 hours. Cell growth or viability was measured using ATPlite, and results were quantified by monitoring luminescence intensity or examining trypan blue incorporation using automated cell counter (Countess II FL, Life Technologies).

### Colony-forming and sphere-forming assays

For colony-forming assays, 500 or 1,000 PDA cells were seeded in a 6-well plate, and wells were monitored daily by microscopy for colony formation. After 10 days, colonies were visualized by crystal violet staining (Sigma). Colony number was analyzed using ImageJ software (NIH). For spheroids formation, 24 hours after indicated transfections, 200 KPC cells in 20 μl medium were seeded in a V-shape 60-well polystyrene mini tray (Nunc). The plate was placed upside down in incubator to allow the formation of hanging drop spheroids. At day 4, spheres were visualized by microscopy and counted.

### Antibodies

The following antibodies were used: RNF125 (Sigma; #HPA041514, western blot dilution 1:1000, immunohistochemistry and immunofluorescence dilution 1:100; Abcam; #ab74373, western blot dilution 1:1000, immunohistochemistry and immunofluorescence dilution 1:100), FLAG (Sigma; #SAB4200071, western blot dilution 1:5000), GAPDH and Tubulin (Santa Cruz; #sc-47724 and #sc-8035 respectively, western blot dilution 1:2500), pan-keratin (Dako; #M3515, IHC dilution 1:100), Ki67(Abcam; #ab15580, IHC dilution 1:100), Cytokeratin-19(CK19, Abcam; #ab52625. IHC dilution 1:100), alpha smooth muscle actin (αSMA, Santa Cruz; #sc-53015, IHC dilution 1:100), PDX1 (Cell Signaling; #5679S, western blot dilution 1:1000, IHC dilution 1:100; R&D Systems; AF2517, IF dilution 1:100), NR5A2 (Invitrogen; #DA5-28347, western blot dilution 1:1000, IHC/IF dilution 1:100; Santa Cruz; #sc-393369, western blot dilution 1:1000), and HDAC2 (Cell Signaling; #2450S, western blot dilution 1:1000, IHC/IF dilution 1:100, ChIP grade #D6S5P). LC3B (#3868S), cleaved caspase-3 (#9661), K63-linkage polyUb (#5621S), K48-linkage polyUb (#4289S), Histone H3 (#3638T), H3K27Ac (#8173S), H3K9Ac (#9649) antibodies were purchased from Cell Signaling and used at 1:1000 dilution for western blot.

### Immunohistochemistry

Sections (5 µM) were cut using a Leica Microsystems cryostat, transferred onto Superfrost-Plus slides (Thermo Fisher Scientific) and stained for H&E. For immunohistochemistry, sections were deparaffinized and rehydrated, and antigen was retrieved using Dako target-retrieval solution (Dako). Endogenous peroxidase activity was quenched by incubation with 3% hydrogen peroxide for 30 min. Specimens were incubated with different antibodies, diluted in Dako antibody diluent overnight at 4°C, washed 3 times with PBS/0.03% Tween-20, and incubated with Dako-labelled polymer-HRP for 1 h at room temperature. Following 3 washes with PBS containing 0.03% Tween-20, sections were incubated with 3,3’-diaminobenzidine chromogen and counterstained with hematoxylin.

### Immunofluorescence

After deparaffinization and antigen retrieval, slides were washed in water followed by PBS, incubated in 0.05% Tween for 15 minutes, and blocked in 10% goat serum (1 hr at room temperature). Slides were then incubated with diluted primary antibody (1:100) in Dako antibody diluent overnight. After 3 PBS washes, slides were incubated with diluted (1:200) secondary antibody for 45 minutes. Following another round of PBS washing, slides were incubated with DAPI for 10 minutes (1:5000 in PBS), washed with PBS, and covered with coverslips in hard mounting medium. Immunofluorescence-stained slides were visualized using a fluorescence microscope with an Aperio slide scanner.

### Immunoprecipitation and immunoblotting

To detect protein interactions, cell lysates were prepared using 1% Triton-lysis buffer (50 mM Tris-HCl [pH 7.4] and 150 mM NaCl, 1 mM EDTA, 1% [v/v] Triton X-100), which included a mixture of protease and phosphatase inhibitors (Halt^TM^, Thermo Scientific). To differentiate cytosolic from nuclear fractions, fractionation was carried out using a subcellular protein fractionation kit (Thermo Scientific Pierce) according to the manufacturer’s instructions. To immunoprecipitate HDAC2 and RNF125, lysates pre-cleared with Protein A/G agarose beads (Santa Cruz) were incubated with indicated antibodies overnight at 4°C. Protein A/G agarose beads were added for an additional 2 hr at 4°C to capture complexes. To immunoprecipitate FLAG-tagged RNF125, lysates were incubated with anti-FLAG M2 affinity beads (Sigma) for 2 hr at 4°C. Beads were washed with lysis buffer, boiled in Laemmli buffer, and subjected to SDS-PAGE. For immunoblotting, cells, organoids, or tissue lysates were prepared using RIPA buffer (50 mM Tris-HCl [pH7.4], 1% [v/v] NP-40, 0.1% [w/v] sodium deoxycholate, 0.1% [w/v] SDS, 150 mM NaCl, 1 mM EDTA, plus a protease inhibitor cocktail ^16^ and PhoStop ^16^). Imaging of immunoblots was performed using a ChemiDoc™ imaging system (BioRad) and respective HRP-conjugated antibodies.

### siRNA and DNA constructs, transfection and transduction

siRNA targeting human RNF125 (#1; SASI_Hs01_00176610, #2; SASI_Hs01_001766112), NR5A2 (#1; SASI_Hs01_00061211, #2; SASI_Hs01_00061214), and PDX1(#1; SASI_Hs01_00108076, #2; SASI_Hs01_00108080) were purchased from Sigma. shRNA-containing plasmids targeting human and mouse RNF125 (Human: #1; TRCN0000004231, #2; TRCN0000004234. Mouse: #1; TRCN0000106401, #2; TRCN0000106404, PDX1 (Human: #1; TRCN0000015652, #2; TRCN0000014648. Mouse: #1; TRCN0000086029, #2; TRCN0000086031), NR5A2 (Human: #1; TRCN0000019656, #2; TRCN0000319345. Mouse: #1; TRCN0000026039, #2; TRCN0000025966), and HDAC2 (Human: #1; TRCN0000004819, #2; TRCN0000004820. Mouse: #1; TRCN0000039397, #2: TRCN0000039395) were purchased from La Jolla Institute for Allergy and Immunology. FLAG-tagged RNF125 wild type (WT) and RING mutant (RM) vectors was previously described^27^. The HDAC2 expression vector was constructed in pLX304-V5 Gateway vector from pDONR223-HDAC2 (DNASU #HsCD00005288). PDX1 (Addgene#114304, Addgene#114305), NR5A2 (DNASU#HsCD443475, Addgene#28093) and RFP-tagged histone H2B (Addgene#26001) expressing plasmids were obtained from Addgene or DNASU as indicated. Cells (2.5 × 10^5^ per well) were cultured overnight in six-well plates and transiently transfected using JetPrime transfection reagent (Polyplus Transfection) according to the manufacturer’s instructions. For viral transduction, packaged viral constructs were harvested from media of 293T cells 3 days following their transfection (calcium phosphate) with the plasmids encoding viral envelope proteins and the viral core construct. Target cells were infected with virus particles by spinoculation (1,600 × *g* for 30 min at room temperature) in the presence of 4 μg/ml polybrene (Sigma). Stable clones were established by culturing cells in media containing puromycin (1 μg/ml, InvivoGen) or blastocidin (5μg/ml, Gibco).

### Differential gene expression analysis

To monitor patient survival we followed up the level of UBL expression in specimens from pancreatic cancer patients, using data from the Oncolnc browser (http://oncolnc.org). TCGA samples were analyzed according to survival with logrank analysis. PDA tumor and adjacent normal pancreas mRNA raw expression profiles were downloaded from GEO [accession#: GSE16515].

### RT-qPCR analysis

RNA was extracted from cell lines and homogenized tumors using a GenElute Mammalian Total RNA Purification Kit (Sigma) according to standard protocols. RNA concentration was measured using a NanoDrop spectrophotometer (ThermoFisher). cDNA was synthesized from aliquots of 1μg total RNA using a high-capacity cDNA synthesis kit (Applied Biosystems). Quantitative PCR was performed with SYBR Green I dye master mix (BioRad) and a CFX connect Real-Time PCR System (Bio-Rad). Primer sequences are listed in Table S1. Primer efficiency was measured in preliminary experiments, and amplification specificity was confirmed by dissociation curve analysis.

### Mass spectrometry

Mia PaCa-2 cells were transfected with control plasmid (pcDNA3.0 with a FLAG-tagged C terminus) or plasmid encoding FLAG-tagged RNF125RM. Cells were lysed 24 hr after transfection using 1% Triton-lysis buffer (50 mM Tris-HCl [pH 7.4]) and 150 mM NaCl, 1 mM EDTA, 1% [v/v] Triton X-100, a protease inhibitor cocktail (Phostop, Roche) and pre-cleared with protein A/G agarose beads (Santa Cruz). IP was performed using FLAG-M2-agarose beads (Sigma). Beads were then resuspended with 8M urea, 50 mM ammonium bicarbonate, and processed for LC-MS/MS analysis using a nanoACQUITY system (Waters) coupled to an Orbitrap Fusion Lumos mass spectrometer (Thermo Fisher Scientific) as previously described^45^. All mass spectra were analyzed with MaxQuant software version 1.5.5.1. MS/MS spectra were searched against the *Homo sapiens* Uniprot protein sequence database (version January 2018) and GPM cRAP sequences (commonly known protein contaminants). Post-search data analysis was performed with R-64 bit version 3.5.1, including the R Bioconductor packages such as limma and MSstats.

### Chromatin immunoprecipitation assay (ChIP)

KPC cells (4 × 10^6^ per immunoprecipitation) were fixed in 1% formaldehyde in PBS for 10 minutes at room temperature and then incubated with 0.125 M glycine for 5 minutes. Cells were washed with PBS, incubated in lysis buffer (50 mM Tris-HCl, pH 8.0, 1% SDS, 10 mM EDTA), and sonicated on ice to shear the DNA into approximately 500-bp fragments. The sonicate was centrifuged, and the supernatant was precleared by incubation with protein A/G beads (Santa Cruz Biotechnology Inc.) on a rotating platform at 4°C for 1 hour. Control IgG or appropriate primary antibody was added and incubated overnight at 4°C. Protein A/G was added, and the mixture incubated at 4°C for 2 hours. Beads were washed sequentially with low-salt buffer (20 mM Tris-HCl, pH 8.0, 0.1% SDS, 1% Triton X-100, 2 mM EDTA, 150 mM NaCl), high-salt buffer (as for low salt except 500 mM NaCl), and LiCl wash buffer (10 mM Tris-HCl, pH 8.0, 0.25 M LiCl, 1% NP-40, 1% sodium deoxycholate, 1 mM EDTA). Chromatin was eluted in 120 µl elution buffer (1% SDS, 100 mM NaHCO3) for 15 minutes at 30°C. Samples were centrifuged, and the supernatant was mixed with 4.8 µl of 5 M NaCl and incubated overnight at 65°C. RNase A (2 µl of 10 mg/ml) and proteinase K (2 µl of 20 mg/ml) were added, and the sample was incubated with gentle shaking 1 hour at 45°C and then extracted with phenol/chloroform to isolate DNA. Samples were subjected to qPCR to detect the PDX1 promoter region, with RPL30 (#7015 Cell Signaling) serving as internal control, using primers shown in table S1.

### Statistics and reproducibility

Statistical significance between two groups was assessed by an unpaired Student’s *t-*test. Ordinary one-way analysis of variance (ANOVA) was used to analyze more than two groups. Two-way ANOVA was used to analyze cell proliferation at multiple timepoints. GraphPad Prism 7 and 8 software (GraphPad) were used for all statistical calculations. All cell culture experiments were performed 3 times, except for the MS candidates screened for effect on PDX1 (Fig. 6B), which was performed 2 times. Data are presented as means ± SD (as noted in figure legends), and *P* values <0.05 were considered statistically significant.

## Supplementary Figure Legends

**Fig. S1.**
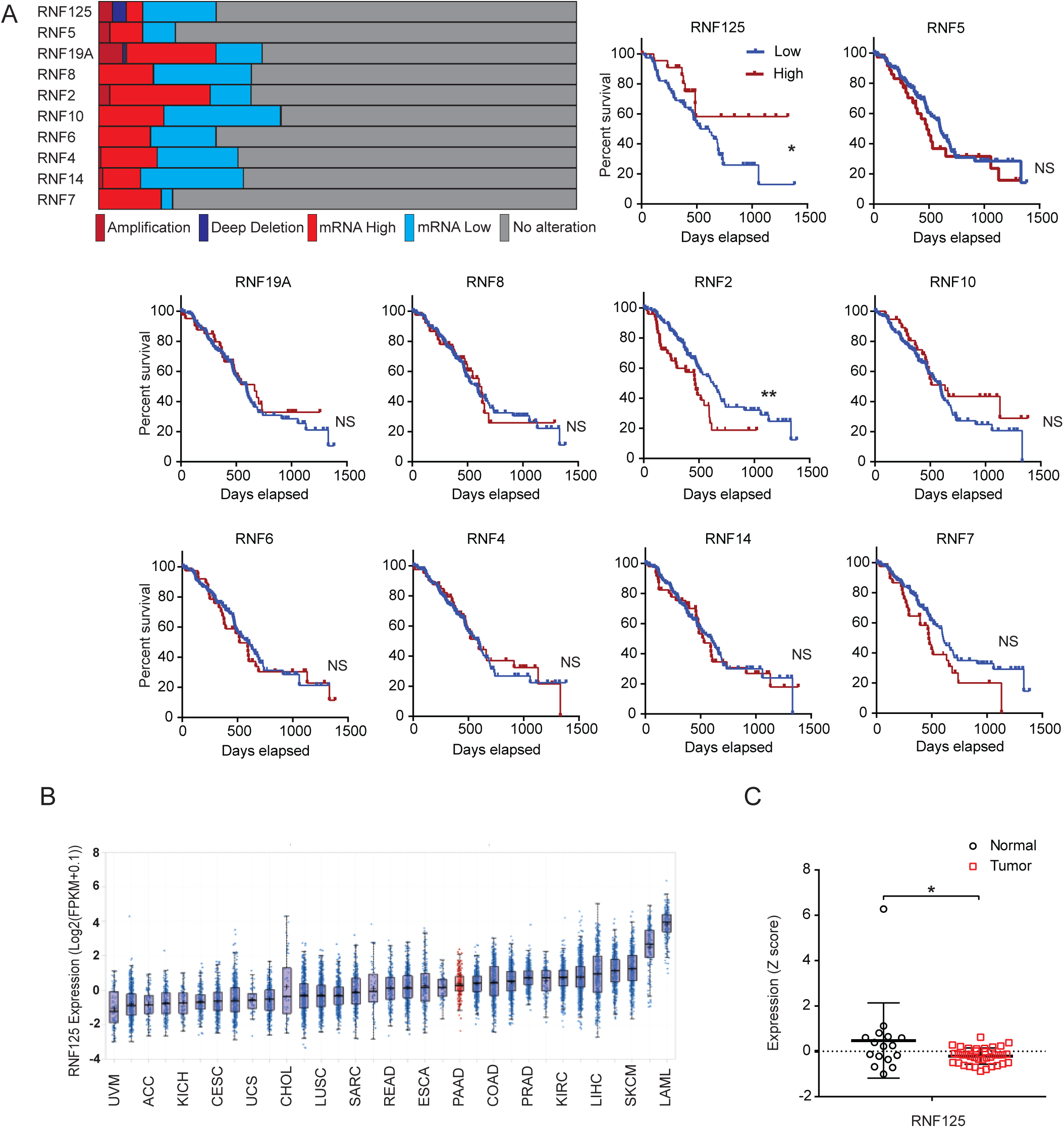
High RNF125 expression in pancreatic cancer is positively correlated with prolonged survival. (**A**) Analysis of TCGA dataset in terms of gene copy number and expression of RNF125 and other genes encoding RING finger proteins in cancer, and effects on patient survival. Kaplan-Meier analysis and log-rank test were used. (**B**) RNF125 expression in different human cancers as analyzed in the TCGA dataset. (**C**) RNF125 expression in normal human pancreas (*n*=16) and in PDA specimens (*n*=36) based on the GSE16515 dataset. Student’s t-test was used to calculate statistical significance. \**P*<0.05, \*\**P*<0.01, NS – nonsignificant.

**Fig. S2.**
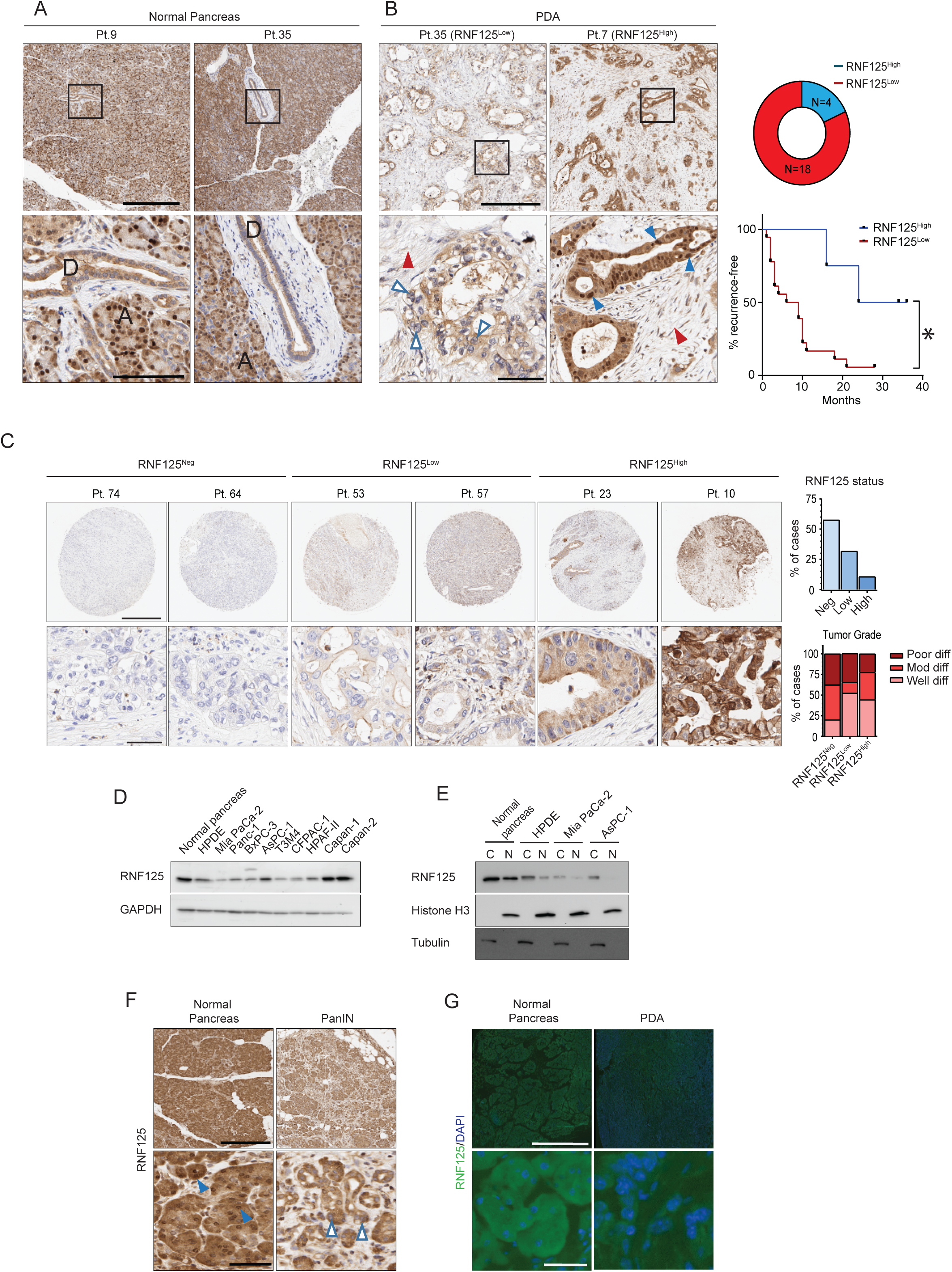
RNF125 level and localization in normal pancreas and in pancreatic cancer. (**A**) RNF125 staining in two representative samples of normal human pancreas tissue. Acinar (A) and ductal (D) structures are indicated. Scale bar = 500 μm (upper) and 100 μm (insets). (**B**) left, RNF125 staining in two samples of human PDA in RNF125^Low^ (left) and RNF125^High^ (right) subgroups. Scale bar, 500μm (upper) and 100μm (insets). Filled arrowheads mark positively stained nuclei, empty arrowheads mark negatively stained nuclei. Red arrowheads mark the negatively stained stromal component of the tumor. right, Pie chart shows numbers of cases per subgroup (upper) and survival in subgroups (lower). (**C**) left, RNF125 staining in representative cases from a pancreatic cancer TMA. right, Graphs depict RNF125 status (upper) and tumor grade as a function of RNF125 levels (lower) across all specimens. (**D**) Western blot analysis of indicated proteins in specimens of human normal pancreas, human pancreatic ductal epithelium (HPDE), and different human PDA cell lines. (**E**) Western blot analysis of indicated proteins in cytoplasmic and nuclear fractions of specimens from murine normal pancreas, human pancreatic ductal epithelium (HPDE), and two human PDA cell lines (Mia PaCa-2, AsPC-1). Histone H3 and tubulin serve as controls for nuclear and cytoplasmic fractions, respectively. (**F**) RNF125 staining in human normal pancreas and PanIN. Scale bar, 200 μm (upper) and 40 μm (inset). Arrowheads mark positive (filled) or negative (empty) nuclei. (**G**) RNF125 staining in normal murine pancreas and a KPC orthotopic tumor. Scale bar, 500 μm (upper) and 50 μm (lower).

**Fig. S3.**
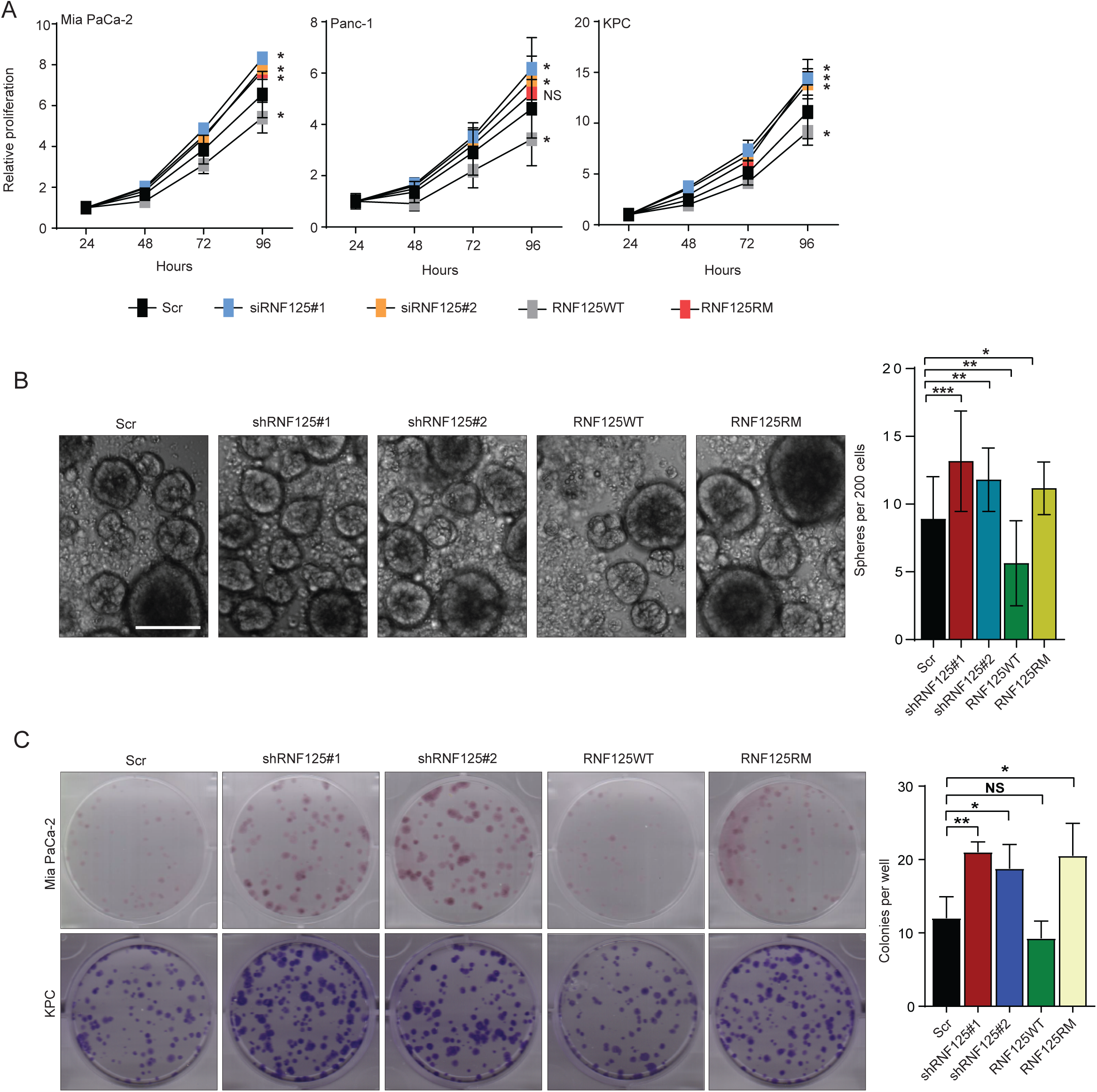
Functional analysis of RNF125 in PDA. (**A**) Growth of three indicated lines upon transfection with Scr, one of two shRNF125s, RNF125WT or RNF125RM. (**B**) Sphere-forming capacity of KPC cells transduced with Scr, one of two shRNF125s, RNF125WT or RNF125RM. KPC cells were seeded 8 hours after transfection, and spheres counted 72 hours later. Scale bar, 50μm. (**C**) left, Plates showing anchorage-dependent colony formation by Mia PaCa-2 or KPC cells transduced with constructs described above. right, Quantification of colony formation assay.

**Fig. S4.**
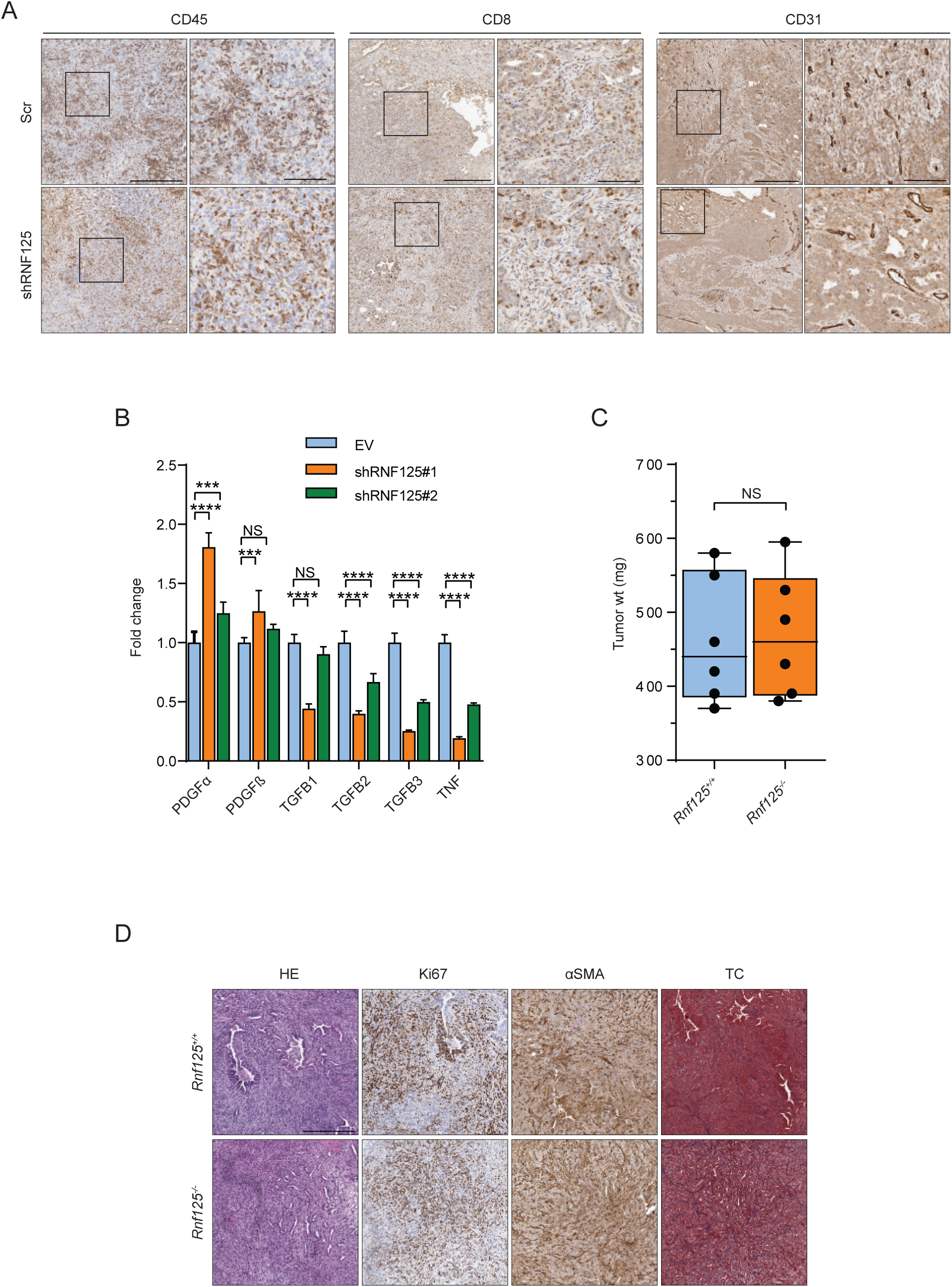
Effects of RNF125 knockdown on the PDA tumor microenvironment. (**A**) Orthotopic KPC tumors transduced with Scr or shRNF125 plasmids and stained for indicated markers. Scale bar, 400 μm (left panels), 100 μm (insets). (**B**) Levels of transcripts encoding fibrogenesis factors in WT and in two shRNF125-transduced KPC tumors. Results are means ± SEM of 2 different tumors per group. (**C**) Weight of orthotopic KPC tumors 3 weeks after injection into the pancreas of WT or *Rnf125^−/−^* mice. (**D**) Indicated staining of KPC tumors grown in WT or *Rnf125^−/−^* mice. TC, Trichrome.

**Fig. S5.**
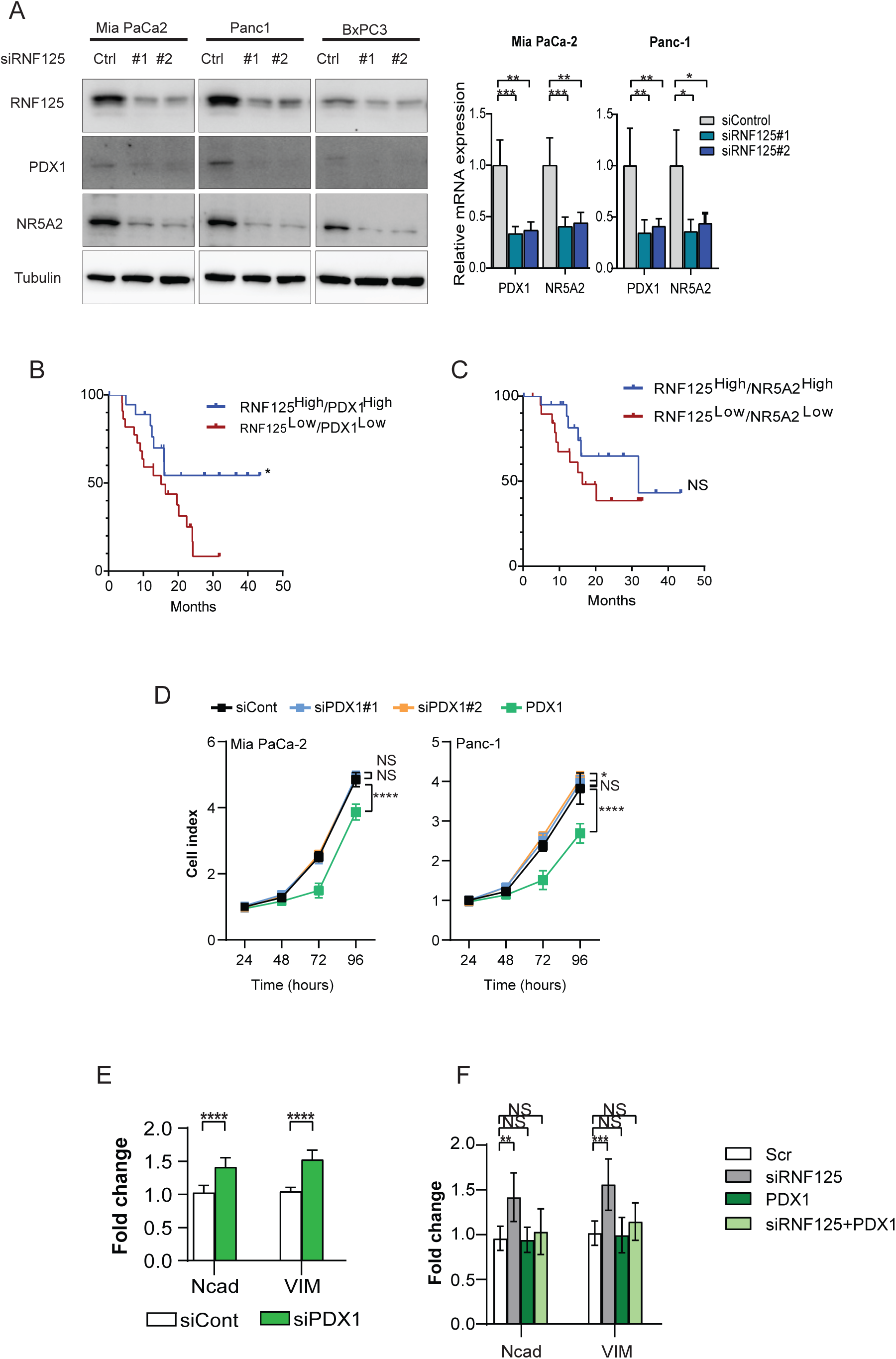
RNF125 - PDX1 in pancreatic cancer. (**A**) Western blot (left) and qPCR (right) analyses of PDX1 and NR5A2 levels after RNF125 knockdown. qPCR results are means ± SEM in 1 of 2 experiments. (**B**) TCGA PDA dataset analysis showing combined effect of PDX1 and RNF125 levels on pancreatic cancer survival and indicating survival of RNF125^High^/PDX1^High^ (*n* = 19) compared to RNF125^Low^/PDX1^Low^ (*n* = 22) patients. Kaplan-Meier analysis and log-rank test were used; two-sided log-rank *P* = 0.0359. (**C**) Similar analysis for effects of NR5A2 levels on pancreatic cancer survival and indicating survival of RNF125^High^/NR5A2^High^ patients (*n* = 19) versus RNF125^Low^/NR5A2^Low^ (*n* = 22) patients. Kaplan-Meier analysis and log-rank test were used. NS – nonsignificant. (**D**) Proliferation of Mia PaCa-2 or Panc-1 lines after knockdown or overexpression of PDX1. Data are presented as means ± SEM of 3 different experiments. Statistical significance was calculated using two-way ANOVA. (**E**) RT-qPCR analysis of transcripts encoding EMT markers in Mia PaCa-2 cells 96 hours after transfection with siPDX1. Data are representative of 3 different experiments with 3 technical replicates. Statistical significance was calculated using two-way ANOVA. (**F**) RT-qPCR analysis of transcripts encoding the EMT markers Ncad or VIM in Mia PaCa-2 cells 96 hours after transfection with indicated constructs. Data are representative of 3 different experiments with 3 technical replicates. Statistical significance was calculated using two-way ANOVA.

**Fig. S6.**
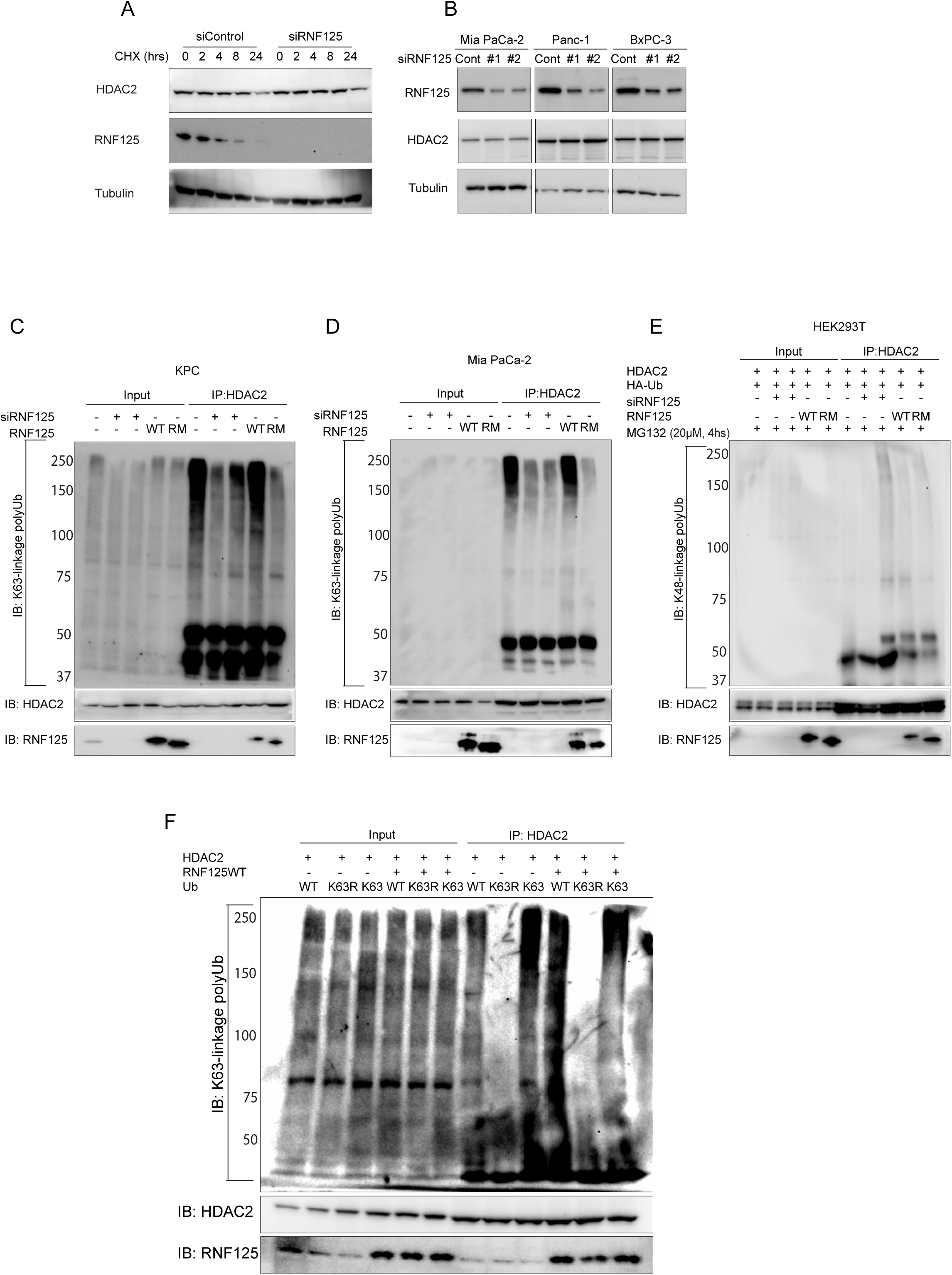
RNF125 interacts with HDAC2 and alters its activity. (**A**) NR5A2 transcript levels in Mia PaCa-2 cells transfected with indicated siRNAs. Results represent <u>mean</u> ± SEM in a representative experiment. (**B**) Western blot analysis of HDAC2 stability in Mia PaCa-2 cells with and without RNF125 knockdown. Seventy-two hours after siRNA transfection, cells were treated with 20μM cycloheximide for indicated times. (**C**) Western blot analysis with indicated antibodies upon RNF125 knockdown in different cell lines. (**D, E**) K63 ubiquitination of endogenous HDAC2 as analyzed in Mia PaCa-2 (D) and KPC (E) cells expressing different levels of RNF125. IP, HDAC2; immunoblot, anti K63-linked Ub antibody. (**F**) K48 ubiquitination of HDAC2 as analyzed in HEK293T cells expressing HDAC2 and HA-Ub, under different RNF125 conditions. IP, HDAC2; immunoblot, anti K48-linked Ub antibody. (**G**) K63 ubiquitination of HDAC2 as analyzed in HEK293T cells expressing HDAC2, HA-Ub, and RNF125, in the presence or absence of K63R. IP, HDAC2; immunoblot, anti K63-linked Ub antibody. \**P*<0.05, \*\**P*<0.01, \*\*\**P*<0.005, \*\*\*\**P*<0.001, NS – nonsignificant.

**Fig. S7.**
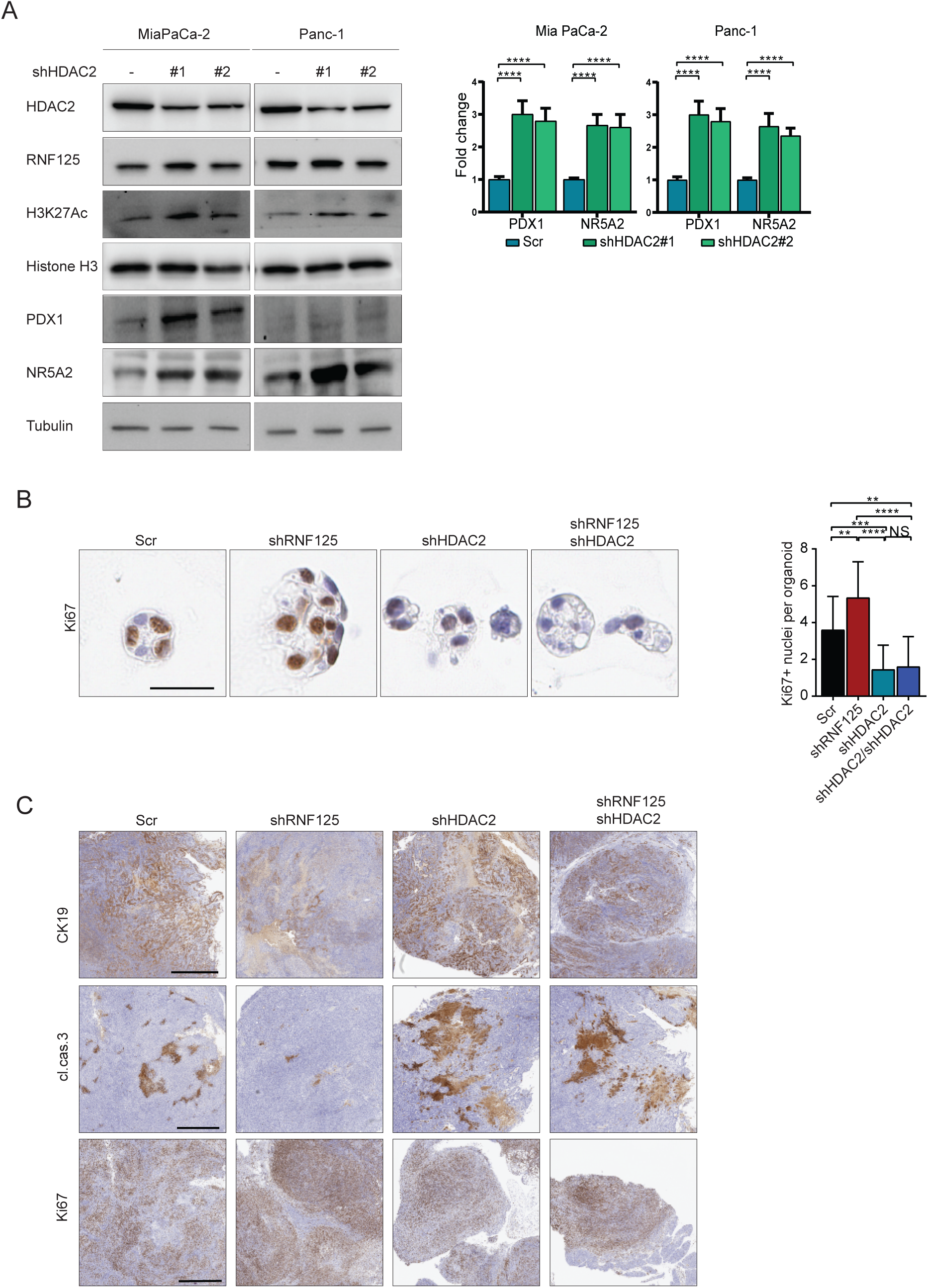
HDAC2 regulates PDX1 and NR5A2 in pancreatic cancer. (**A**) left, Immunoblotting with indicated antibodies and (right) qPCR analysis of PDX1 and NR5A2 transcripts 96 hours after transfection of Mia PaCa-2 or Panc-1 cells with Scr, shHDAC2#1, or shHDAC2#2 plasmids. qPCR results are means ± SEM in 2 independent experiments. Statistical significance was calculated using two-way ANOVA. (**B**) left, Ki67 stain in 48-hour-old KPC organoids transduced with Scr, shRNF125s, shHDAC2, or a shRNF125/shHDAC2 combination. right, Quantification of Ki67-positive nuclei. Statistical significance was calculated using one-way ANOVA. Scale bar, 100µm. (**C**) CK19, cleaved caspase-3, and Ki67 staining of orthotopic KPC tumors, either control or transduced with shRNF125, shHDAC2, or both. \**P*<0.05, \*\**P*<0.01, \*\*\**P*<0.005, \*\*\*\**P*<0.001, NS – nonsignificant

